# Depth-Dependent Advantages of Three-Photon Microscopy for Imaging Through Intact Murine Cortical Bone and Into the Marrow

**DOI:** 10.64898/2026.07.31.742087

**Authors:** Samantha Bratcher, Michael R. E. Lamont, Chris B. Schaffer, Karl J. Lewis

**Affiliations:** Meinig School of Biomedical Engineering, Cornell University, Ithaca, NY, USA

**Keywords:** *In vivo* Imaging, Bone, Three Photon Microscopy, Two Photon Microscopy

## Abstract

Bone adapts to its mechanical and physiological environment through the coordinated actions of osteocytes and marrow-resident cell populations. Because these processes depend on complex interactions within native tissue microenvironments, intravital imaging offers a unique opportunity to reveal cellular behaviors that cannot be fully captured *ex vivo*. However, the highly scattering nature of mineralized bone limits imaging depth and direct observation of cells within intact tissue. Two-photon (2P) microscopy has enabled important advances in bone biology, while three-photon (3P) microscopy has been proven to extend imaging depth and image quality in skeletal tissues. Here, we directly compared the performance of 2P and 3P microscopy in *ex vivo* mouse long bones by quantifying laser attenuation, signal-to-noise ratio, signal-to-background ratio, and spatial resolution, as well as assessed the impact of wave-front correction. We found that 2P and 3P microscopy generated comparable image quality through approximately 50 *µ*m of cortical bone. Beyond this depth, however, 3P microscopy provided superior brightness, contrast, and resolution, enabling improved visualization of structures deep within the cortex and at the cortical–marrow interface. To assess compatibility with intravital imaging, we evaluated endogenous markers of cellular stress during 3P imaging. Although prolonged continuous imaging decreased osteocyte spontaneous calcium signaling magnitude and increased autofluorescence, short intermittent imaging bouts produced negligible evidence of cellular damage compared to controls. Leveraging the enhanced penetration depth of 3P microscopy, we further show visualization of immune cell migration within the marrow cavity through intact cortical bone in both the third metatarsal (MT3) and tibia. Together, these findings affirm 3P microscopy as a powerful tool for studying cellular dynamics in living bone. By extending imaging beyond superficial cortical regions and enabling direct visualization of marrow-resident cells through intact bone, 3P microscopy expands opportunities to investigate osteocyte biology, marrow niche function, and skeletal adaptation in vivo.

## Introduction

Bone is a dynamic tissue that plays important roles beyond structural support and protection. It continually adapts to mechanical stimuli, becoming stronger or weaker in response to physical demands due to the actions of osteocytes, cells which comprise more than 90% of the bone cell population (1–4). These cells reside within bone in the lacunocanalicular network (LCN), a complex system of caves and tunnels, where they sense mechanical strain and regulate bone formation and resorption (5–8). The geometry of this local microenvironment is crucial to osteocyte mechanosensitivity and variations to its morphology may play a role in aging and disease related changes in bone (9–12). In addition to mineralized tissue, bone contains marrow, an essential home for hematopoiesis, immune cell development, mesenchymal stem cells, and support of bone homeostasis (13, 14). Evidence further suggests osteocytes are able to directly impact development and regulation of these marrow cell populations (15, 16).

Despite the importance of studying osteocytes and the marrow within, investigations within fully intact bone can be technically challenging. Bone tissue is optically dense. A high degree of mineralization, low water content, and tight organization of collagen matrix result in significant light scattering and limited optical penetration into bone tissue. Advances in optical imaging have progressively enabled visualization of osteocyte behavior from glass slides to their native microenvironment. Confocal microscopy of osteocytes *in situ* connected fluid-induced shear stress within the LCN to osteocyte response (6, 17). Later, two-photon (2P) microscopy paired with mechanical loading of the mouse MT3 enabled direct visualization *in vivo*, confirming previous in situ findings (5). The effective imaging depth and quality of 2P microscopy remains limited by optical scattering and absorption, particularly in deeper regions such as the endocortical surface and marrow cavity. More advanced microscopy techniques are needed to interrogate deep osteocyte and marrow-resident cell populations *in vivo*.

Three-photon (3P) microscopy has been proven as a powerful tool that offers enhanced depth penetration and improved signal fidelity relative to 2P microscopy, particularly deep below the tissue surface. The advantages of 3P microscopy arise primarily from the higher order nonlinear excitation process, which decreases background signal generation outside of the focal plane, boosting signal relative to the background. Concurrently, the longer wavelengths used for 3P imaging experience less scattering and absorption, allowing 3P microscopy to penetrate farther into tissues (18–20). These properties have enabled 3P microscopy to achieve imaging depths exceeding 1mm in the mouse brain (19). Recent work by Rakhymzhan et al. established the feasibility of 3P imaging through intact bone, demonstrating penetration into the marrow space of intact mouse tibial cortex using excitation wavelengths greater than 1600nm (21). Building upon this advance, important opportunities remain to evaluate the utility of 3P microscopy to visualize fluorescent signal within bone and its functionality with more commonly used green fluorescent reporters.

Although 3P microscopy has been shown to generate quality images deep below bone tissue, a further complication in multiphoton microscopy of whole bone noted by Rakhymzhan et al. is wavefront distortion caused by the introduction of optical aberrations by the curved bone surface (21). Light rays that strike a curved interface are refracted to different angles along the radius of curvature. Consequently, the rays fail to converge to a single focal point, introducing astigmatism into the imaging system. Strategies to mitigate wavefront distortion may attempt to correct the generated image (adaptive optics) or minimize the source of distortion (refractive index matching) (Figure 1). Adaptive optics (AO) employs a deformable mirror with a precisely controlled surface profile to generate opposing aberrations in the laser and effectively cancel distortions. Alternatively, refractive index (RI) matching introduces an RI-matched medium above the curved sample, allowing excitation light to maintain its trajectory without refraction and improving signal quality. Additional implementation of these wavefront correction strategies may grant further benefits to previously reported 3P capabilities.

**Fig. 1.**
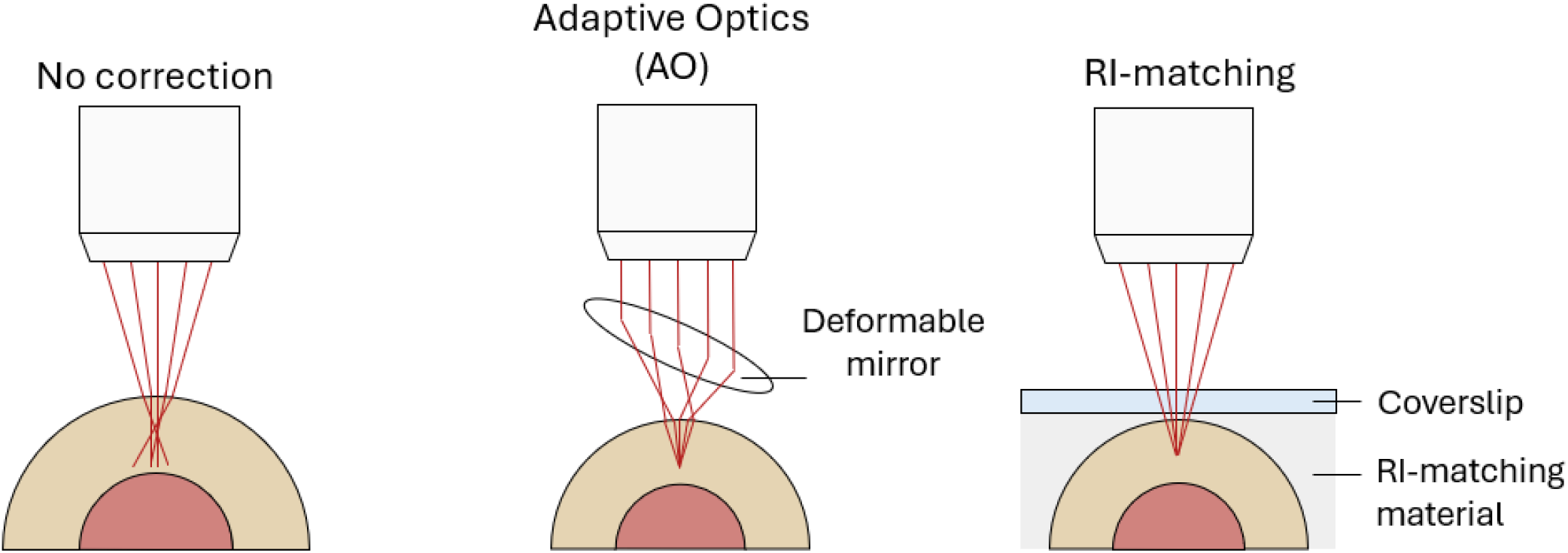
Simplified visualization of wavefront distortion caused by the curved surface of bone and correction methods. (Left) Curved surfaces introduce unequal degrees of refraction, preventing light from converging to a single focal point and degrading image quality. (Middle) AO pre-empts distortions within the tissue by introducing distortions in the laser before tissue penetration, effectively canceling distortions. (Right) RI-matching materials applied above the bone surface provide a smoother optical transition, changing the interface from curved to flat.

The demonstrated performance of 3P microscopy to penetrate bone motivates further exploration of its capability within bone when utilizing common fluorescent markers along with wavefront correction strategies. In this study, we quantitatively compared the laser penetration and image quality between 2P and 3P microscopy in mouse long bones and assessed the extent to which wavefront correction improved imaging performance. We hypothesized that 3P imaging yields superior depth-dependent signal preservation, normalized signal intensity, signal-to-noise ratio, and signal-to-background ratio compared to 2P, with further enhancement achieved through AO and RI-matching materials. Additionally, we evaluated whether 3P imaging can be applied to bone *in vivo* over extended periods without inducing signs of cellular damage and furthermore exemplified the use of 3P for functional observation into the marrow cavity of intact cortical bone.

## Methods

All procedures were approved by the Cornell University IACUC (Protocol #2020-0046).

### Microscope Setup

All images were collected using either a custom 2P or 3P microscope. Two-photon laser excitation was achieved using a Chameleon laser (Coherent) at 920nm excitation. Three-photon laser excitation was achieved using an Opera OPA (Coherent) pumped by a Monaco laser (Coherent) at 1320nm excitation. The ScanImage (MBF Bioscience) module in MATLAB (MathWorks, Version R2022b) was used to control microscope laser power and image collection, which used a 25x objective (Olympus, XWMP) with an NA of 1.05. A bandpass filter of 540/80nmwas used to isolate fluorescent signal. Surface laser power did not exceed 110mW to avoid tissue damage.

### Ex vivo image collection

Mice with osteocyte-targeted expression of GCaMP8m were created by crossing DMP1-Cre mice (B6N.FVB-Tg(Dmp1-cre)1Jqfe/BwdJ; JAX Labs, Strain#023047) with mice containing GCaMP8m knocked-in under the Igs7 locus (STOCK Igs7tm1(tetO-GCaMP8m,CAG-tTA2)Genie/J; JAX Labs, Strain#037718) (22, 23). GCaMP8m served as a fixable fluorescent marker of osteocytes within bone. The third metatarsals (MT3), tibias, femurs, and humeri of skeletally mature (16-18-week-old) females were dissected, fixed in zinc-buffered formalin (Anatech Ltd) for 24-48 hours, and soaked overnight in 1 mM CaCl_2_ to enhance calcium indicator fluorescence. A landmark was created 1mm proximal to the mid-diaphysis for consistent imaging location across bones.

Wavefront correction was achieved using a refractive index (RI) matching adhesive, adaptive optics (AO), or a combination of both. RI matching was implemented during 2P and 3P imaging by applying Loctite 4305 adhesive (Henkel) to the bone surface followed by placement of a 5mm round glass coverslip. This approach was chosen based on published success imaging through mouse calvaria (24). Light pressure was applied to remove excess glue from between the bone and coverslip. The adhesive was allowed to air-dry for 5 minutes before curing under UV light. AO was implemented during 3P imaging with a deformable mirror (Bertin ALPAO, D69) controlled by a custom MATLAB (Mathworks, Version R2022b) app. The mirror surface was manually defined to maximize overall image brightness using the first fourteen Zernike polynomials. AO was incorporated in 3P alone as aberrations are expected to more prominently affect 3P, where fluorescence scales with the cube of the focal intensity. Thus, modest reductions in intensity due to aberrations can produce substantial signal loss. Furthermore, the objective of this study was to evaluate the enhance performance of an optimized 3P imaging modality, rather than compare AO performance in 2P.

To assess signal decay caused by laser attenuation through bone, z-stacks were acquired starting at the surface and moving deeper into the tissue at 5*µ*m intervals while maintaining constant laser power. In a separate experiment to evaluate image quality with depth, z-stacks were collected from the surface at 2*µ*m intervals with exponentially increasing laser power. All z-stacks continued below the surface until fluorescent signal was no longer distinguishable. Average surface laser power was 60mW and did not maximally exceed 110 mW to avoid tissue damage.

### Quantification of laser penetration and image quality

Image stacks acquired with constant laser power through depth were cropped in FIJI (NIH) to the center 100*µ*m to remove uneven fluorescent intensity across the field-of-view caused by the curved bone surface (Figure 2). Cropped image stacks were then imported into MATLAB for calculation of effective attenuation length (EAL). At each depth, the mean intensity of the top 1% brightest pixels was calculated, and the natural log of the signal was plotted as a function of depth. The depth corresponding to the peak signal intensity was identified, and a linear regression was performed from this point through the linear decay region of the curve. EAL was defined as the depth at which the natural log of the signal intensity decreased by 1/e^2^ (2P) or 1/e^3^ (3P) according to equation 1 or 2 respectively.

**Fig. 2.**
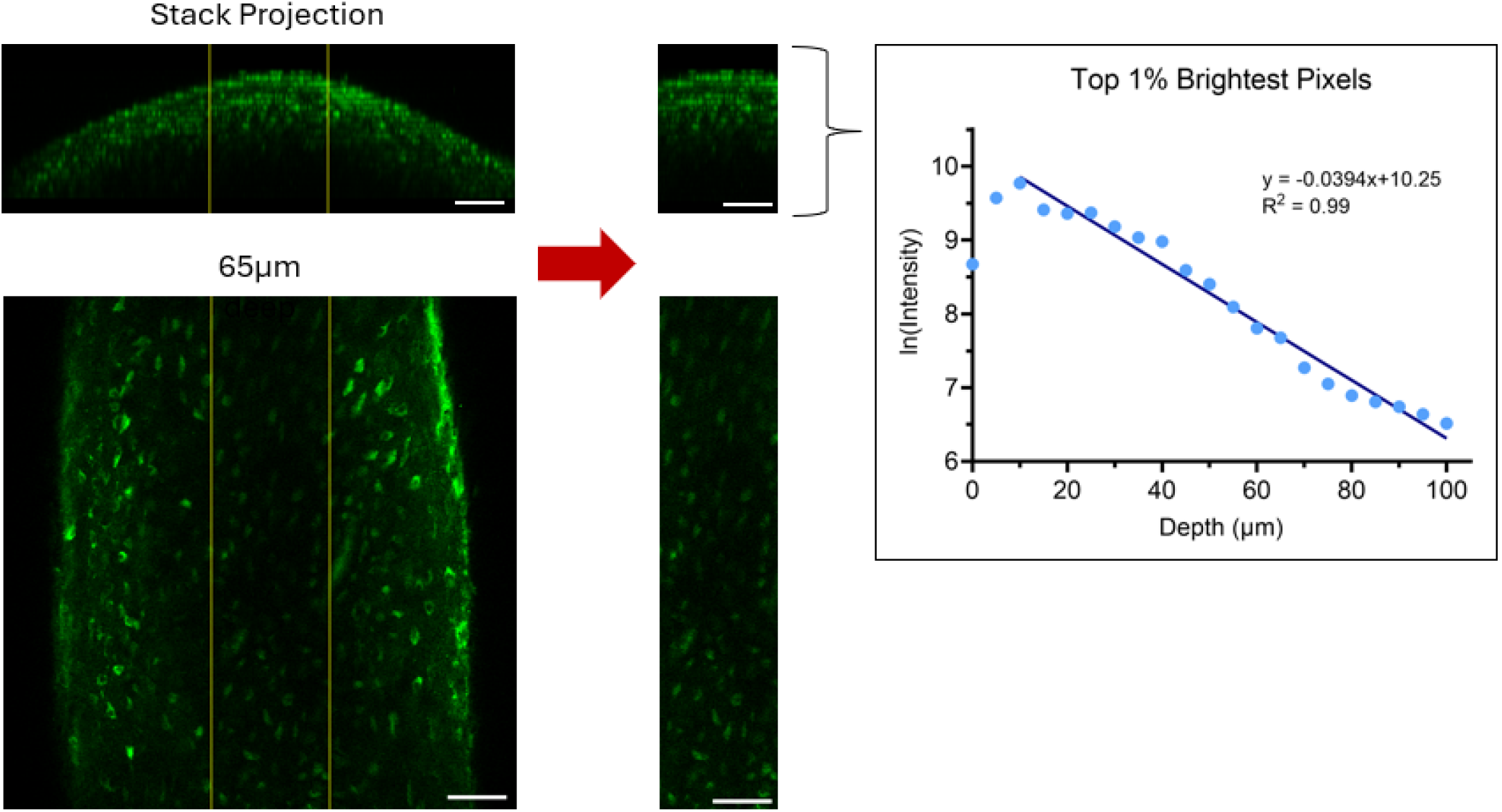
Quantitative workflow for calculation of EAL with a representative image from a mouse humerus. Image stacks were first cropped to the center 100*µ*m (outlined in yellow) to remove uneven fluorescent intensity across the field-of-view. Cropping is important, as this effect is more noticeable as depth increases, example shown at 65*µ*m below the surface. After cropping, the mean intensity of the top 1% brightest pixels was calculated, and the natural log of the signal was plotted as a function of depth. Scale bar = 50*µ*m.

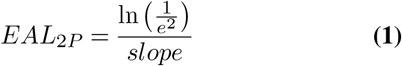

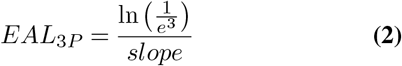

Image stacks collected with exponentially increasing power over depth were also cropped in FIJI (NIH) to the center 100*µ*m. Cell regions of interest (ROIs) were then generated for quantification of image quality. An auto-local threshold was used to generate a binary image of cells followed by morphological operations (open, close, fill) to refine segmentation. ROIs were identified using automatic particle counting with a size range of 50-180 pixels^2^. Raw mean intensity of each ROI was measured for every z-slice. Background ROIs for intensity normalization were manually created at 10*µ*m intervals. All ROI intensity values were imported into MATLAB (Mathworks, Version R2021a). Background intensity values were interpolated across slices, and cell signal intensity was normalized accordingly. Signal-to-noise ratio (SNR) was calculated as the average raw cell intensity divided by the average background standard deviation. Signal-to-background ratio (SBR) was calculated as the average raw cell intensity divided by the average background intensity.

Image resolution was determined by manual selection of cells at the peak and bottom of the image stack. A line was drawn across the edge of the cell and background in FIJI (NIH). Intensity values across the line were imported to MATLAB and fit against an error function according to equation 3.

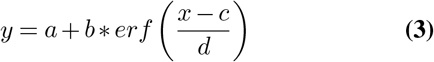

### Simulation of adaptive optics correction through a curved surface

To better understand the key optical aberrations affecting whole bone imaging, optical simulation was performed in Zemax OpticStudio (Version 2025 R1.00, Ansys, Inc.) in sequential mode. The entrance pupil diameter was matched to the optimized fill fraction for the back aperture of the objective used for the experiments (25). To model the effects of the deformable mirror, the first surface was defined as a Zernike standard phase type, which applies a phase described by Zernike polynomials from astigmatism to quadrafoil (Z5 to Z15 in Noll ordering) with prescribed amplitudes. The objective was then modelled simply as a paraxial surface with a focal length matching the effective focal length of the experimental objective. This was followed by a standard surface with a refractive index of 1.32, corresponding to water at room temperature and a 1320nm wavelength (26). The final surface represented the bone and was modelled using an extended polynomial along the x-axis fitted to the measured average surface profile of each sample with a refractive index of 1.56 (27). Simulations were performed centered at the apex of the bone curvature and offset laterally ±30*µ*m.

The Zemax system was controlled programmatically from MATLAB (Mathworks, Version R2024b) via Zemax’s Interactive Extension, enabling easy setting of the parameters, trigger optimization, and extraction of resulting data. For each bone type, imaging depth, and lateral offset, the focus was optimized by varying both the water thickness (distance between the objective lens and bone surface) and Zernike polynomial amplitudes with a metric based on the wavefront referenced to the chief (central) ray. Following optimization, the resulting point spread function, *I*(*x, y*), calculated using the Fast Fourier Transform method, was used to determine the effective area at the focus according to equation 4. This was repeated with all Zernike polynomial amplitudes fixed at zero, varying only the water thickness, to attain a focal effective area without adaptive optics.

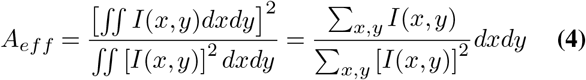

### In vivo estimation of cell stress in response to laser exposure

The application of 3P microscopy in living tissues raises important concerns regarding cellular and tissue health. Risk of phototoxicity or thermal damage from absorption of the excitation laser are expected to increase with continuous imaging time. To assess cell stress associated with laser exposure, mouse MT3s were subjected to imaging while osteocytes were monitored for changes in spontaneous calcium signaling and accumulation of autofluorescent proteins. Osteocytes display spontaneous calcium signals under static conditions, and numerous proteins involved in cellular metabolism, such as NADH, are intrinsically autofluorescent (28–30). Alterations in either of these signals may be suggestive of cell stress in response to imaging.

Skeletally mature (23-24 week old) female DMP1-Cre;GCaMP6f mice were anesthetized with 3% isoflurane and maintained at 1.5% throughout the procedure. The MT3 was surgically isolated for intravital imaging as previously described (5). Briefly, an incision was made above the MT3, overlying tendons resected, and a pin inserted between the MT3 and underlying tissue to isolate it. The hindpaw with pin was positioned in a custom holder within a room-temperature DPBS bath and secured with a bracket arm.

We evaluated two paradigms of laser exposure designed to mimic either brief, intermittent observation of gradual changes to cellular activity or prolonged continuous imaging of dynamic cellular activity. Three-photon excitation at 1320nm was focused 30*µ*m below the bone surface. Intermittent exposure consisted of repeated 3-minute illumination periods separated by 12-minute rest intervals for a total duration of 75 minutes (Figure 3). In contrast, continuous exposure extended to 15 minutes to a total duration of 75 minutes. MT3s that did not receive any additional laser exposure beyond that required for timepoint collection served as procedural controls to assess the effect of surgery alone. A final set of controls were generated by disarticulating the hind paw immediately prior to imaging to induce tissue hypoxia and establish the baseline change due to injury. This group will be referred to as a hypoxic reference. Cell stress measurements were taken every 15 minutes for 75 minutes. At each time-point, a 150s time series (256×256 pixels, 4.22fps, 1.5 zoom, 1320nm excitation) was collected to record spontaneous calcium signaling followed by a 5 frame time series (512×512 pixels) using 820nm excitation to capture autofluorescence of proteins used in cellular metabolism. To reduce animal use, both MT3s from each mouse were used with starting group randomized. Laser power was maintained at 10mW at the bone surface for all conditions.

**Fig. 3.**
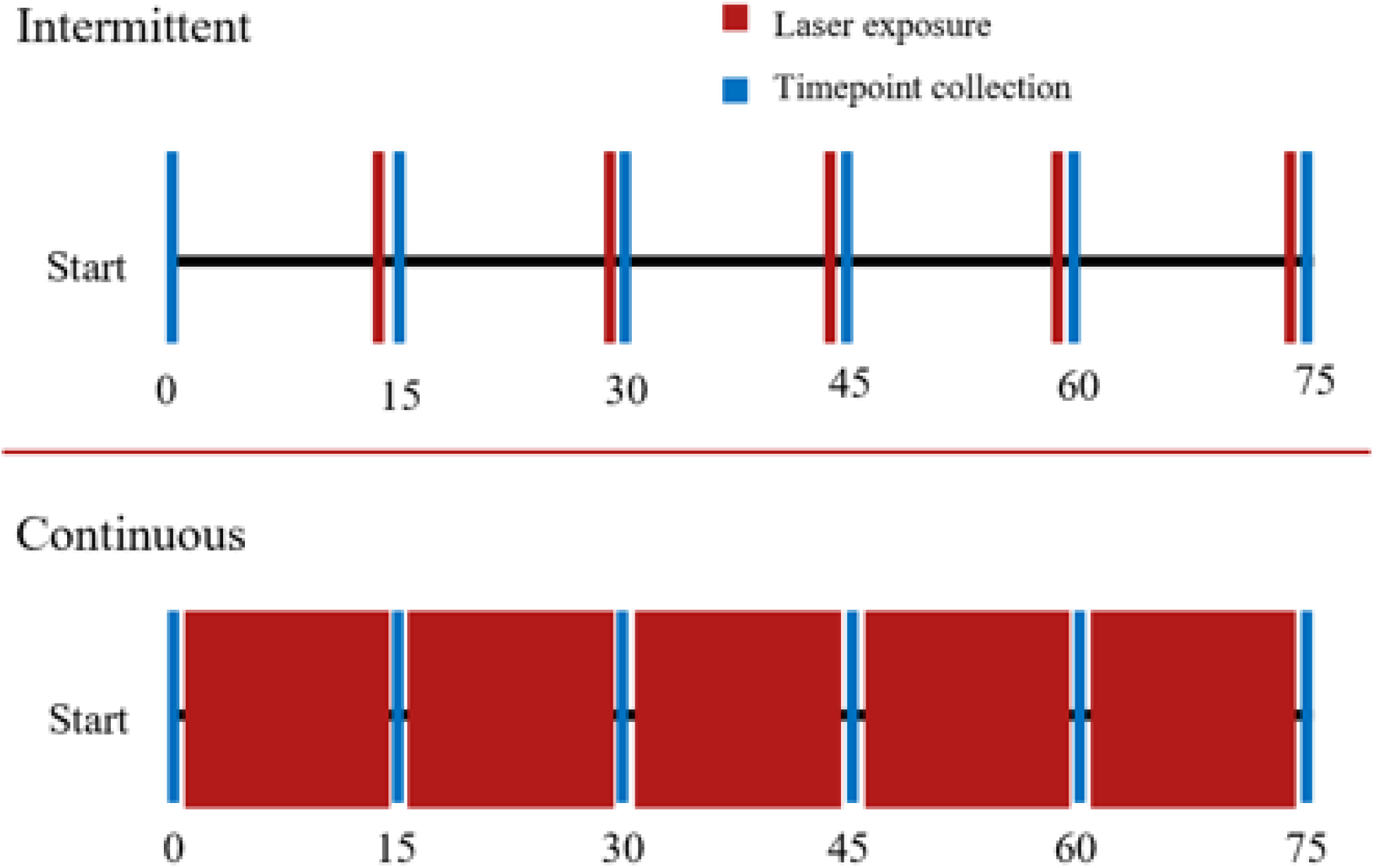
Timeline of data collection and laser exposure to monitor osteocyte cell stress during intravital imaging. (Top) Intermittent groups were exposed to the laser in 3 minute bursts with 12 minutes of rest between exposure bouts. (Bottom) Continuous groups were exposed to the laser for a continued 15 minutes. In both groups, a timepoint was collected every 15 minutes.

### Quantification of in vivo cell stress indicators

Time series of osteocyte calcium signaling were aligned with template matching in FIJI (NIH). Osteocyte ROIs were generated using a manual threshold and automatic particle analysis with a size range of 25-400 pixels^2^. Raw intensity values of cells over time were imported into MATLAB (MathWorks, Version R2021a), normalized to the background, and corrected for photobleaching using a linear detrend. Cell ROIs with an average intensity lower than three standard deviations above the background were excluded as out-of-plane cells. Remaining cell intensities were passed through a low-pass filter. Spikes in ROI intensity were identified as peaks based on a prominence greater than 0.01. Spike magnitudes were calculated as a percentage difference between the static and loading portion of the time series per each cell. Actively signaling cells had a greater than 25% increase in peak intensity during loading.

Images containing potential cell stress induced autofluorescence at each time point were concatenated into a stack in FIJI (NIH) and aligned using linear stack alignment with scale-invariant feature transform (SIFT) based on GCaMP signal. An average projection through time was created, and cell ROIs were segmented by user defined thresholding, morphological operations, and automatic particle analysis excluding elements less than 50 pixels^2^. ROIs were manually deleted or added as necessary, and three background ROIs selected. Generated ROIs were then used to extract raw intensity changes over time in the autofluorescence channel. Intensity values were normalized to the background in MATLAB. ROIs with an average intensity lower than three standard deviations above the background were excluded. The SBR calculated as the ratio of raw intensity to the background.

### Observation of immune cell migration in the marrow cavity

A 16-week-old female mouse expressing EGFP in monocytes under the CX_3_CR1 gene was used (B6.129P2(Cg)-*Cx3cr1*^*tm1Litt*^/J; JAX Labs, Strain #005582) (31). The mouse was anesthetized using 3% isoflurane and maintained at 1.5% throughout the procedure. First, the third metatarsal (MT3) was surgically exposed and isolated as previously described (5). Briefly, an incision was made over the MT3 and overlying tendons removed. A pin was placed between the MT3 and underlying tissues, isolating it. The MT3 was then placed in a room temperature PBS bath to maintain hydration, and a bracket placed over the pin secured the MT3 in place. The objective was focused on the mid-diaphysis, and a z-stack was acquired from the periosteal surface into the marrow space. An additional time series was recorded just below the endocortical surface, identified by collagen second harmonic generation fluorescence, to observe immune cell migration. Images were taken every 30 seconds for 15 minutes. Following MT3 imaging, the mouse was euthanized via cervical dislocation. The tibia was dissected and placed immediately into PBS. Z-stacks and time series were repeated at the anterior surface of the metaphysis.

### Statistical analysis

Significance between EAL values was determined by two-way ANOVA with multiple comparisons between imaging modality and surface condition. Image quality metrics were compared first by linear regression from the peak to the bottom. Regression slopes were compared by one-way ANOVA with multiple comparisons. Osteocyte calcium signaling and laser induced autofluorescence were compared with two-way ANOVA or a mixed model with multiple comparisons. All statistics were performed in Prism (GraphPad, Version 9) (p < 0.05).

## Results

### Excitation laser penetrates bone farther with 3P than 2P and is not impacted by wavefront correction

Effective attenuation length (EAL) quantifies the loss of laser power arising from scattering and absorption of a material. By measuring fluorescent signal intensity as a function of imaging depth under constant excitation power, we observed modality-dependent differences, with brighter cells in three-photon microscopy (3P) and 3P with adaptive optics (3P+AO) compared to 2P (Figure 4). Quantification of the brightest 1% of pixels confirmed EAL is greater using 3P than 2P in all bones tested (i.e. MT3, tibia, femur, and humerus), indicating 3P excitation penetrates bone material more effectively than 2P (Figure 5). Neither wavefront correction via AO nor RI-matching glue affected EAL (Figure 6).

**Fig. 4.**
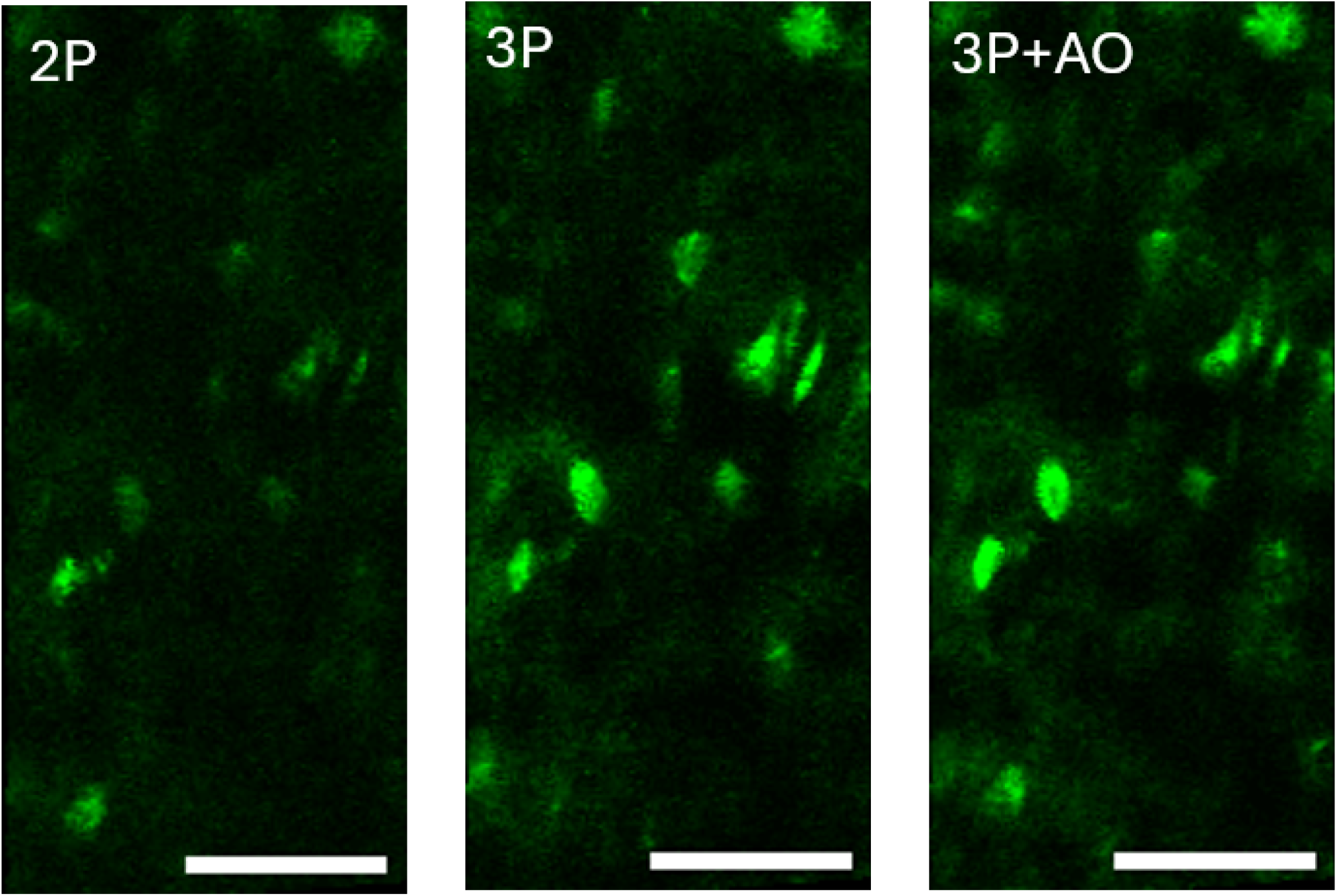
Representative images collected using 2P microscopy, 3P microscopy, and 3P microscopy with AO in a humerus 45*µ*m below the surface. Osteocyte fluorescent signal is distinctly brighter using 3P and 3P+AO than 2P. Scale bar = 50*µ*m.

**Fig. 5.**
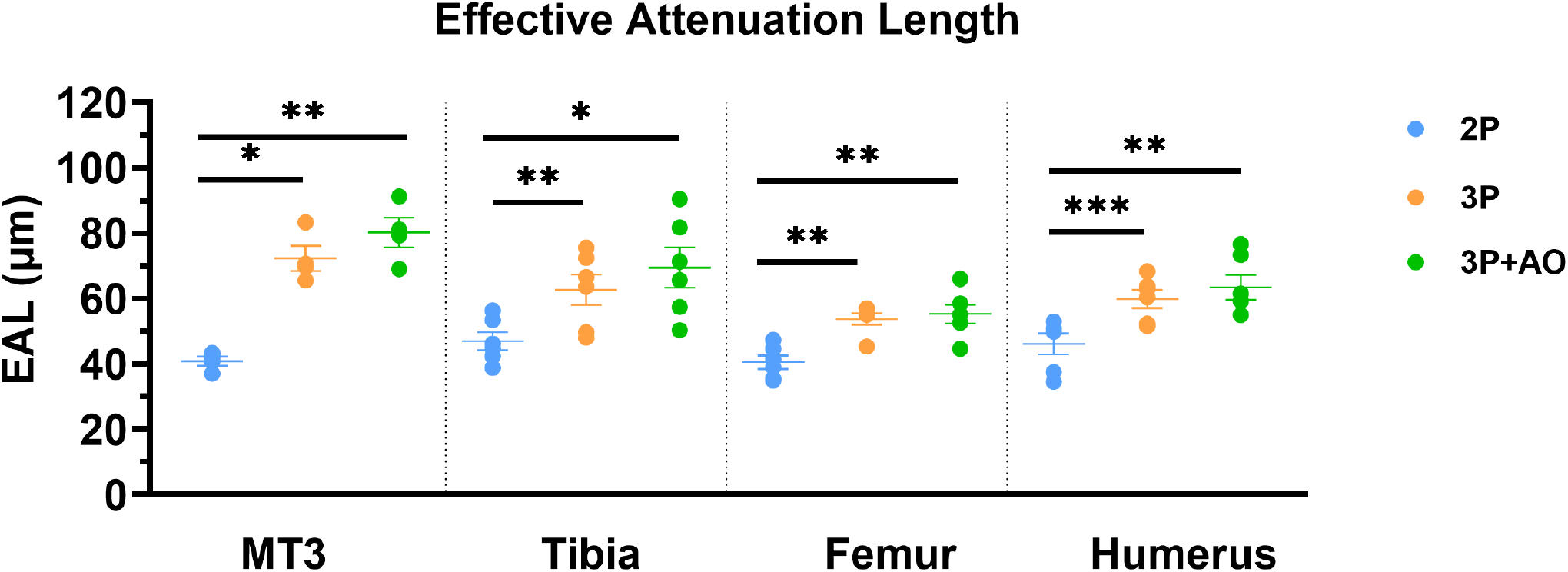
Effective attenuation length (EAL) in the four long bones examined *ex vivo*. EAL is longer using 3P than 2P in all bones tested, indicating more effective material penetration with 3P. Interestingly, AO did not affect EAL in 3P image stacks.

**Fig. 6.**
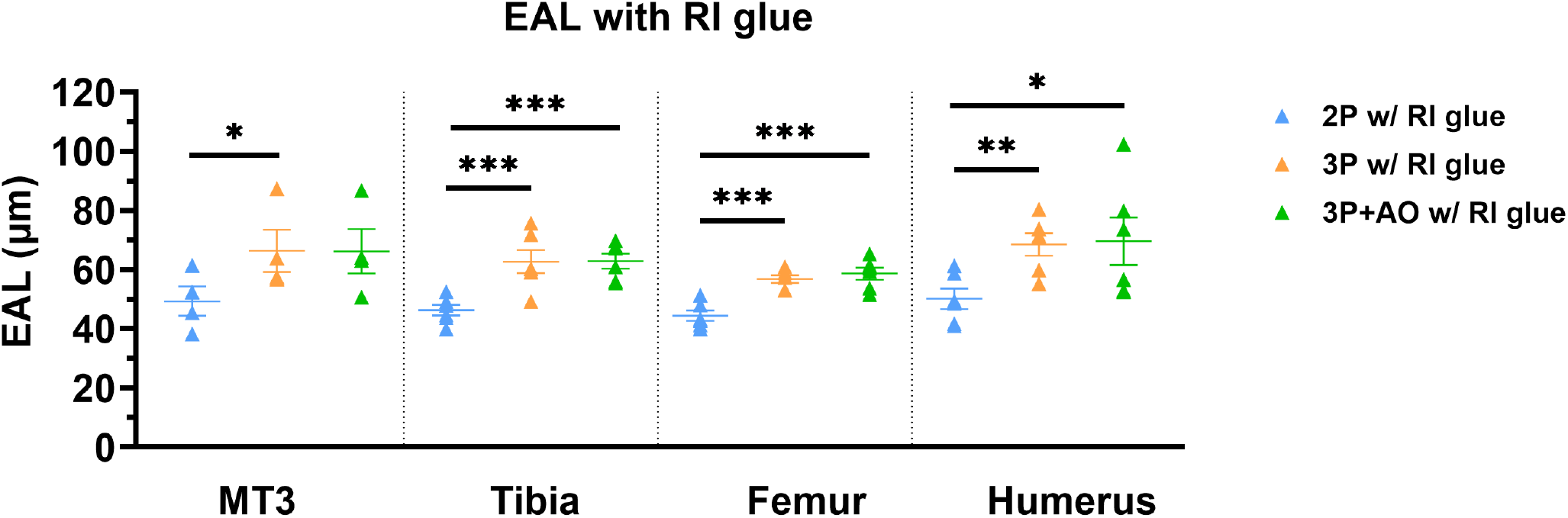
EAL in the four long bones examined with the addition of RI-matched adhesive to the bone surface. EAL values were unchanged, with 3P maintaining a longer EAL than 2P.

### Three-photon microscopy produces brighter images with better resolution

To further build a more complete understanding of imaging performance across depth, we assessed the influence of imaging modality and wavefront correction on the detection of a fluorescent marker within osteocytes as a function of depth in bone. Signal-to-background ratio (SBR) is one useful indicator of signal brightness relative to the background and may serve as a proxy for image quality. In the tibia and humerus, 2P delivered higher initial values that decreased rapidly below 3P and 3P+AO beyond ~40-50*µ*m (Figure 7). In contrast, in the MT3 and femur, 2P SBR was initially comparable to 3P and 3P+AO but remained lower as depth increased. The addition of AO with 3P did not perform better than 3P alone. Adding RI-glue increased the peak SBR values across conditions. In the MT3, SBR decay was increased with RI-glue but values themselves were substantially larger. In the femur, RI-glue simply slowed 2P decay rate. Notably, a crossover in 2P SBR was noticed in two bones; RI-glue extended the depth at which 2P exceeded 3P by ~10*µ*m in the tibia and a surprising ~40*µ*m in the humerus. We observed similar modest, bone-dependent variations in the normal intensity and SNR of fluorescent signal (Supplemental Figures S1 and S2).

**Fig. 7.**
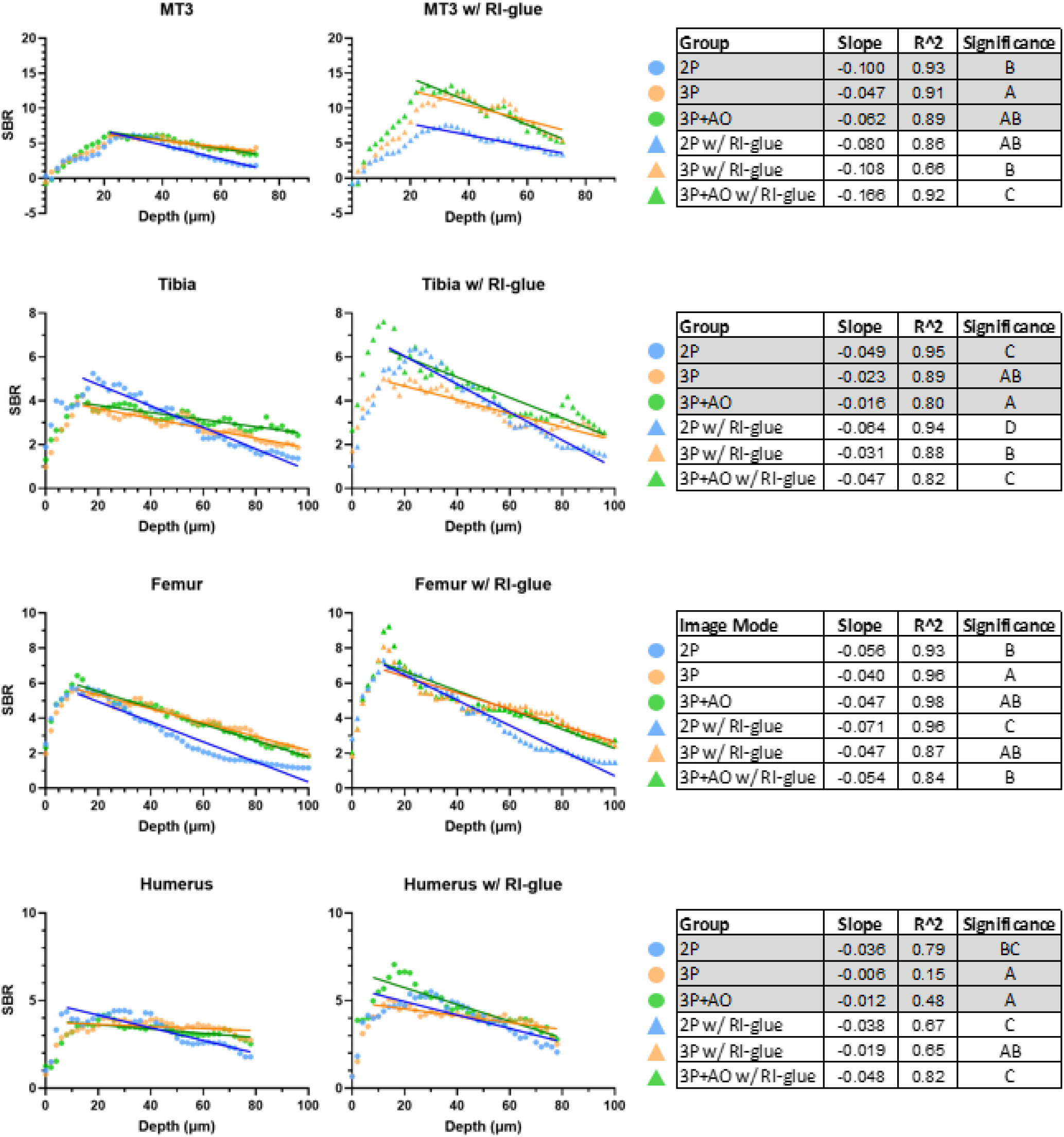
Signal-to-background ratio (SBR) of osteocyte GCaMP8m signal over depth in 2P and 3P microscopy. Large variation in SBR was observed across bones. The tibia and humerus SBR trends were similar to SNR, in which 2P was greater than 3P and 3P+AO until a crossover depth was reached. The MT3 saw dramatic increases to SNR with an increase in 3P and 3P+AO decay, while the femur saw only a modest decrease in 2P decay rate.

Image quality can also be characterized by resolution, or the degree to which an object can be distinguished from its background. Resolution of osteocytes deep below the surface was better in 3P images and further improved with the addition of RI-matching glue (Figure 8). Signal intensity relative to immediate background is notably brighter in 3P and 3P+AO images, further confirmed in quantification of the intensity change from the background across the cell edge. After adding RI-glue, the difference in signal intensity was substantially increased across all conditions, with the cell edge becoming more distinct as shown by a steeper slope of the fitted error function (Figure 8). In addition, 2P signal intensity was improved to levels comparable to 3P+AO images.

**Fig. 8.**
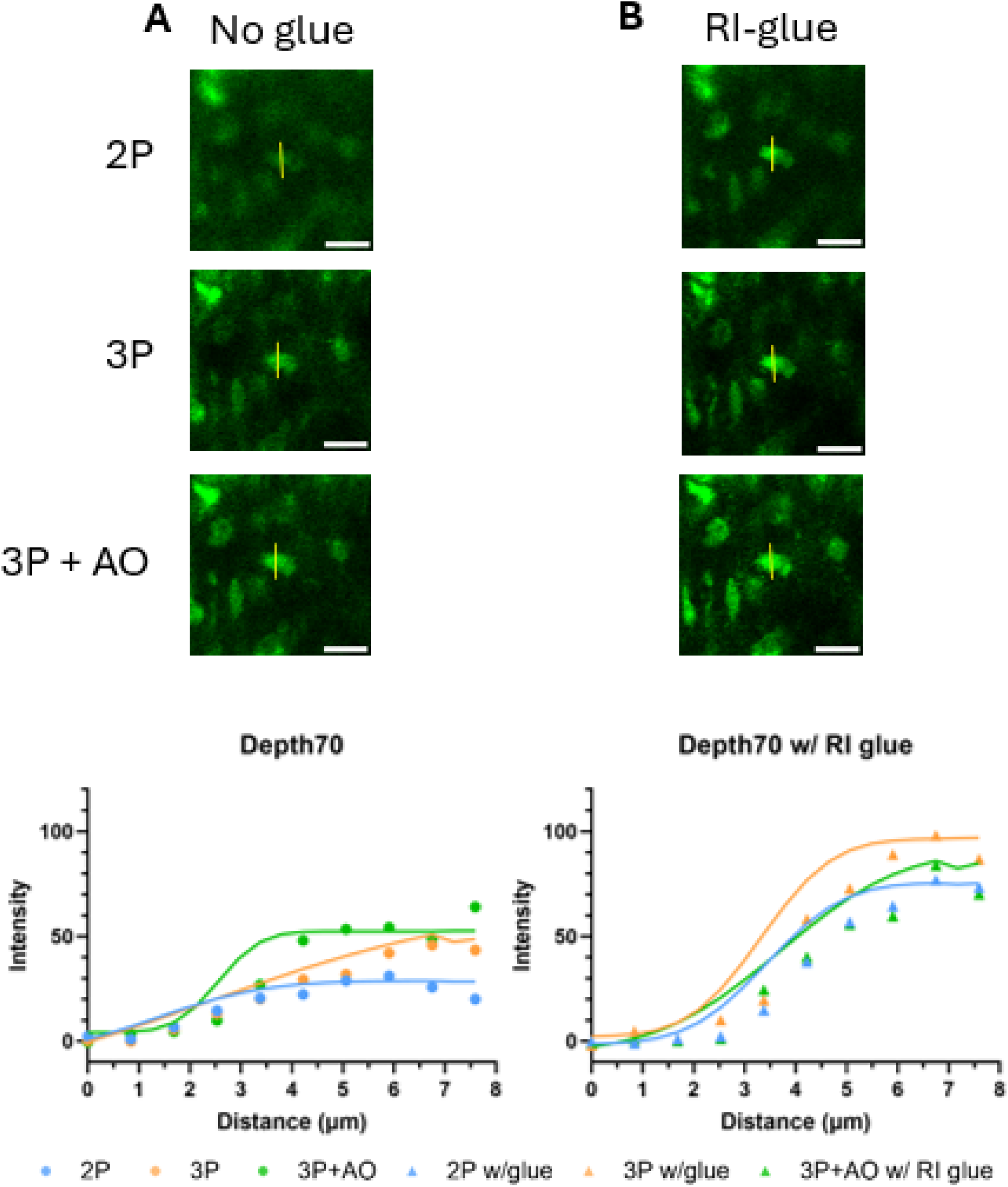
Resolution of osteocytes deep below the bone surface. Representative images in mouse femur cortical bone at 70*µ*m below the surface (A) without and (B) with RI-matching glue. The resolution of osteocytes deep below the surface is greater in 3P and 3P+AO images and further improved by adding RI-matched glue. Examining the intensity across the cell edge highlights the contrast between the cell and background, with steeper slopes indicating a more prominent cell. Scale bar = 20*µ*m

Simulation of the optimal correction with AO identified primary astigmatism as the dominant aberration over imaging depth, with only a minor effect from the primary spherical (Supplemental Table 1). In addition, a greater degree of correction was required with increasing imaging depth; moreover, bones with greater surface curvature required larger correction amplitudes. Simulation further demonstrated AO effectively maintained the beam area throughout the imaging depth (Supplemental Figure S3). In contrast, without AO correction, the beam area increased with depth, with this effect again progressively more pronounced in bones with higher surface curvature. In the MT3, the beam area increased by a factor of 2.3 at a depth of 100*µ*m below the surface, whereas the beam area was nearly the same in the femur. Little variation was observed when off-target from the apex of the surface curve (data not shown).

### Intermittent intravital imaging with 3P is not associated with changes to in vivo markers suggestive of cellular stress

Laser-induced heating and phototoxicity during intravital imaging generate concerns over damage to tissues and cells as imaging duration increases. To assess potential damage caused by the laser over time, we imaged the same region at the mid-diaphysis of the mouse MT3 over 75 minutes using either intermittent or continuous laser exposure conditions. Osteocytes have been observed to display spontaneous calcium signaling under static conditions (28). Time-lapse image series confirmed a small subset of osteocytes generate spontaneous calcium signals (Supplemental Video S1). Osteocytes exposed to the laser intermittently in 3 minute sessions displayed an initial jump in percentage of spontaneously active cells which decreased back to baseline over time (Figure 9). A similar pattern was observed in negative control samples which did not receive laser exposure beyond image collection. Osteocytes exposed continuously for 75 minutes showed an initial high percentage of response which varied over time. Finally, in positive control samples where the hindpaw was disarticulated a consistent low percentage of cells exhibited calcium responses. Across all groups, the percentage of cells determined as responsive remained low at less than 8% throughout the duration of the experiment. Additionally, spontaneously signaling osteocytes produced weak calcium signals when normalized to an initial 40 second time frame. The ratio of response magnitude remained less than 1.5 across all groups.

**Fig. 9.**
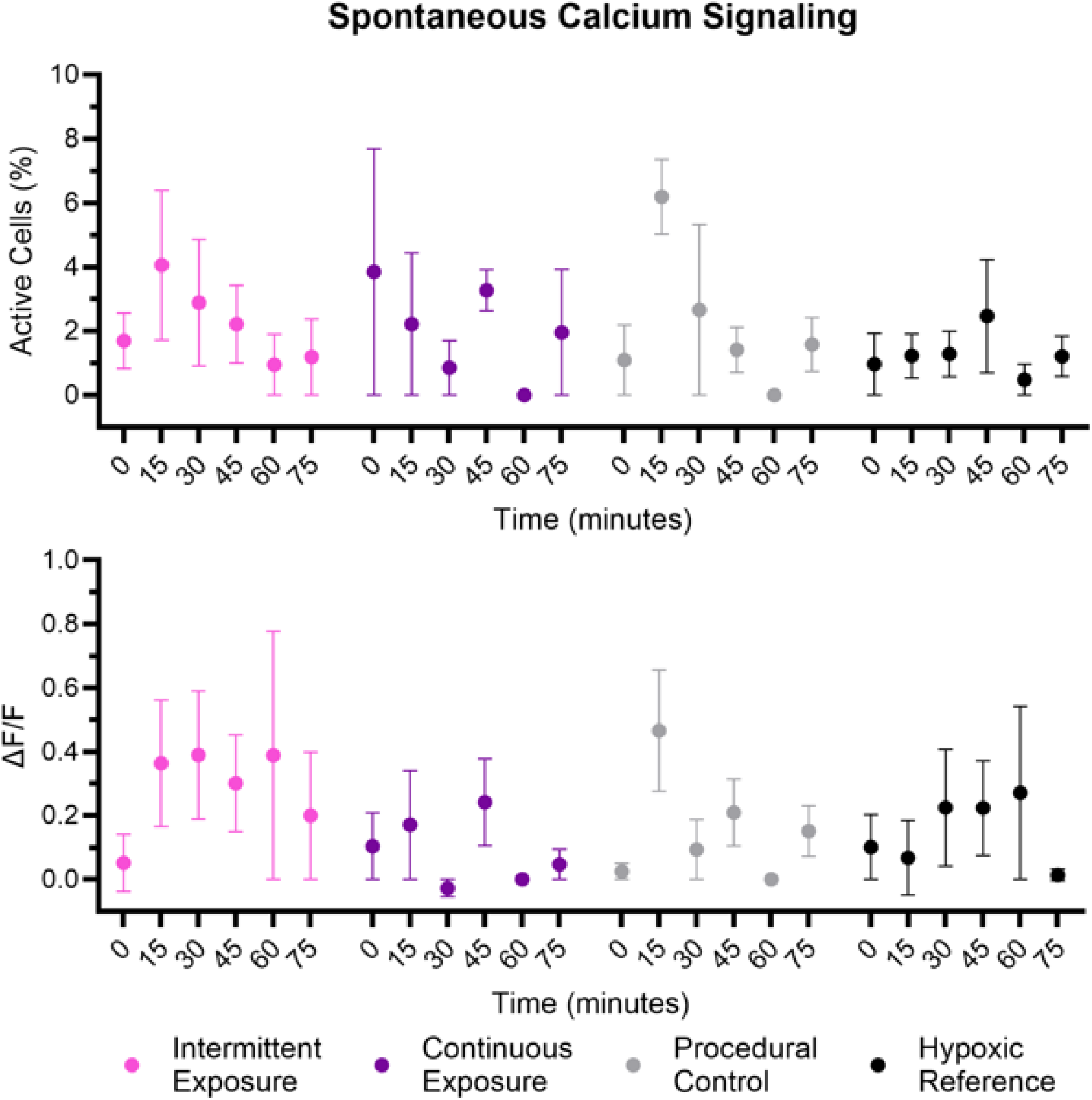
Spontaneous calcium signaling of osteocytes *in vivo* in response to intermittent or continuous laser exposure. (Top) Intermittent, continuous, and negative groups had a sharp increase in percentage of spontaneously active cells within 0-15 minutes, whereas the positive group remained relatively constant in comparison. High variance was observed in the percentage of active cells. (Bottom) Cells that did respond only produced weak calcium signals.

The accumulation of autofluorescent metabolic proteins such as NADH have been correlated with reductions in mitochondrial activity in osteocytes and may serve as another potential indicator of cell stress (29, 30). Cell ROIs with an intensity greater than three standard deviations above the background were classified as positive for autofluorescent signal, and the accumulation of which was calculated by the intensity relative to the background. The percentage of positive cells showed a marked increase over time, as well as an increase in autofluorescent signal (Figure 10). Intermittent, continuous, and negative groups had more positively marked cells than the positive control group, with continuous showing the greatest percentage of positive cells (Figure 10). In positive and intermittent control samples, fluorescent intensity of positive cells increased steadily over time. Both continuous and negative groups increased up to ~45 minutes after initial exposure, after which fluorescent signal varied. The SBR of accumulated autofluorescence increased in all groups over time with the brightest cells generated in the continuous laser exposure group. Although, a change to decreasing signal was noted at 45-60 minutes in the continuous and negative groups.

**Fig. 10.**
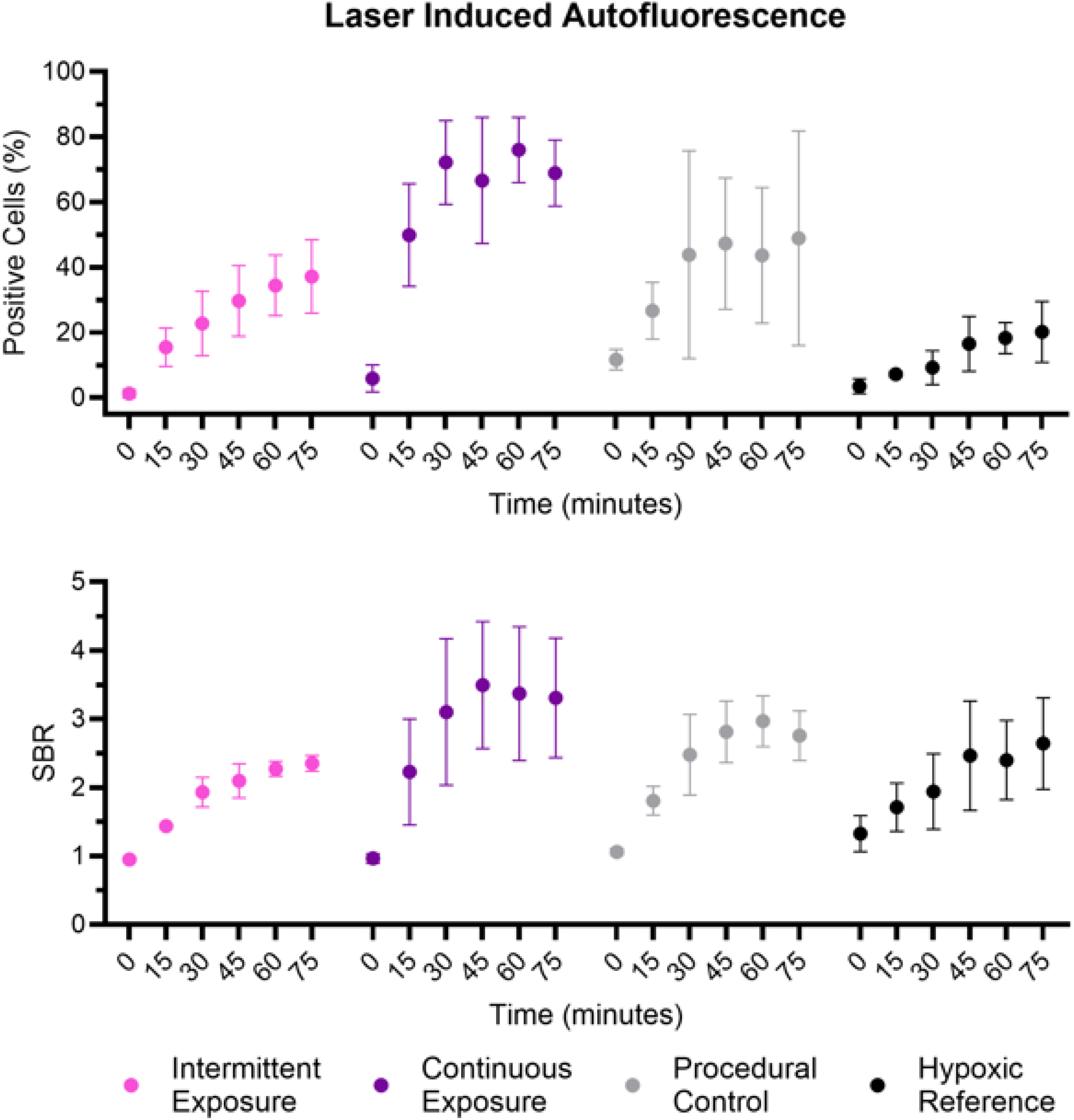
Quantification of laser induced autofluorescence in osteocytes *in vivo*. (Top) Laser exposure markedly increased the quantity of cells denoted positive over time in intermittent, continuous, and negative groups. (Middle) The normal intensity of positive cells slowly increased over time in intermittent and positive groups, while continuous and negative groups reached a peak and began to decrease. (Bottom) The SBR of accumulated signal mirrored the normal intensity, though continuous exposure resulted in greater autofluroescent SBR. Error = SEM.

### Three-photon microscopy penetrates into the marrow space of mouse long bones, granting observation of immune cell migration

Bone marrow is central for continued formation of blood, immune, and bone cells (13, 14). The improved image quality achieved with 3P microscopy at depth enables *in vivo* interrogation of marrow, overcoming the longstanding challenge of imaging through mineralized bone. Using mice with fluorescently labeled immune cells, we visualized structures extending from the bone surface and into the marrow cavity, simultaneously exploiting endogenous harmonic generation signals.

Within the mid-diaphysis of the cortical bone of the MT3, SHG highlighted the collagen matrix, with osteocyte lacunae appearing as dark voids (Figure 11). In parallel, THG revealed the dense network of the LCN. The marrow cavity was reached approximately 100*µ*m below the bone surface, at which point the SHG signal became very diffuse, and the THG signal was replaced by large, closely packed, round cells. Based on morphology and location, THG-positive cells are most likely adipocytes, consistent with the presence of yellow marrow in the long bones of the extremities and previously reported THG-based observations of adipocytes (32– 34). EGFP-positive immune cells were observed interspersed among adipocytes with a small degree of autofluoresence around the periphery of the adipose cells. Notably, we were able to continue capturing marrow and immune cells another 50*µ*m below the endocortical surface into the marrow space, demonstrating the ability of 3P microscopy to probe cellular features deep within the marrow cavity as well.

**Fig. 11.**
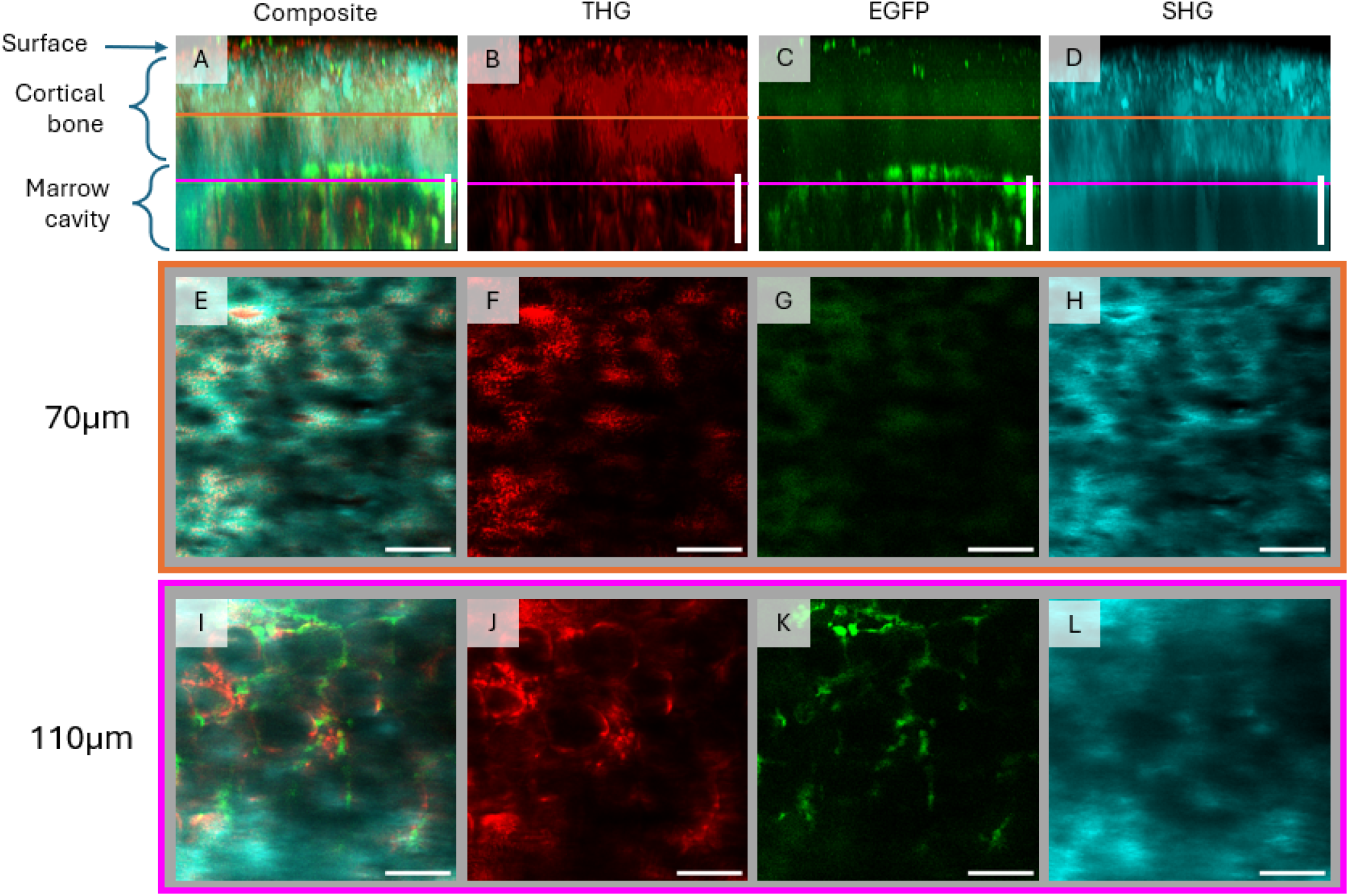
Representative images in the MT3 from the surface into the marrow cavity using 3P. (A-D) 3D projections highlight imaging penetration from the surface into the marrow space. (E-H) 2D image slice from within the cortical bone 70*µ*m below the surface. Lacunae and canaliculi are brightly indicated by THG, whereas canaliculi appear as dark voids in SHG. (I-L) 2D image slice from within the marrow space 110*µ*m below the surface. THG signal gives way to large adipose cells. Immune cells labeled with EGFP are visible between the adipose cells. Scale bar = 50*µ*m.

Successful access to the marrow cavity was further demonstrated through functional imaging of immune cell migration. Time-lapse imaging acquired every 30 seconds over15 minutes captured active movement of EGFP-labeled immune cells within the MT3 marrow cavity (Supplemental Video S2). These cells migrated between the larger adipose cells at varying rates (Figure 12).

**Fig. 12.**
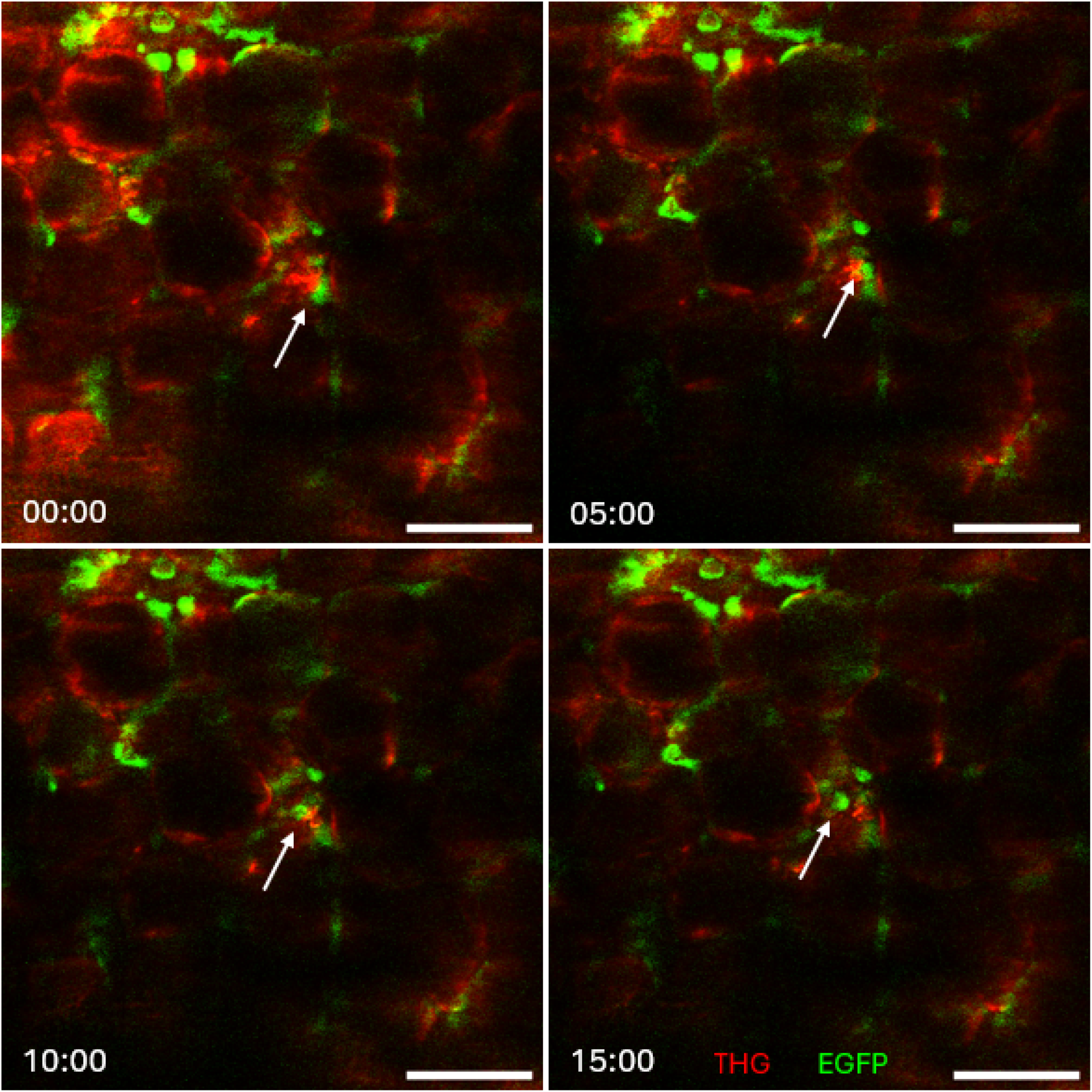
*In vivo* images over 15 minutes in the MT3 marrow cavity using 3P microscopy. Time series images highlight the migration of immune cells (green) between adipose cells (red). A particularly active cell was noticed in the imaging plane shifting location (white arrow). Scale bar = 50*µ*m.

While easily accessed using our lab’s approach, the MT3 midshaft primarily contains yellow marrow, a type primarily functional in energy storage (35, 36). In contrast, red marrow is highly vascularized and suited for hematopoiesis and immune cell formation (14, 37). To further investigate 3P functionality within a differing marrow environment, we also performed in situ imaging of the mouse tibia metaphysis. The cortical bone was consistent with that observed in the MT3, albeit, thicker. The tibia marrow cavity was located approximately 140*µ*m below the surface and contained a much sparser degree of weakly labeled immune cells (Figure 13). THG signal revealed a distinct population of small, clustered cells with small degrees of overlap with EGFP signal. Likewise in the MT3, cells remained detectable for an additional 50*µ*m into the marrow cavity.

**Fig. 13.**
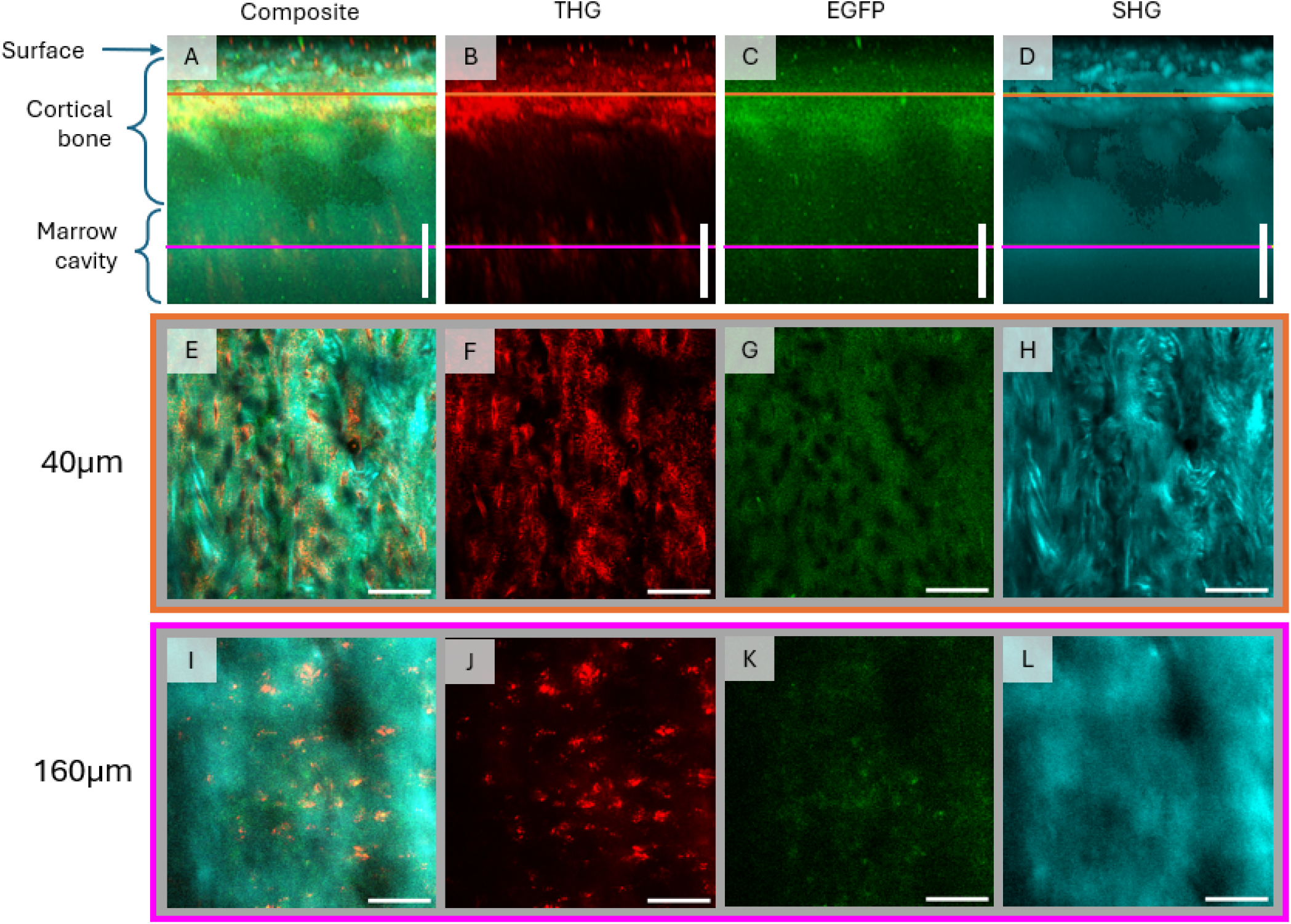
Representative images in the tibia from the surface into the marrow cavity using 3P. (A-D) 3D projections highlight imaging penetration from the surface into the marrow space. (E-H) 2D image slice from within the cortical bone 40*µ*m below the surface. Lacunae and canaliculi are brightly indicated by THG, whereas canaliculi appear as dark voids in SHG. (I-L) 2D image slice from within the marrow space 160*µ*m below the surface. THG signal gives way instead to small, clustered groups of cells. Immune cells labeled with EGFP are weakly visible. Scale bar = 50*µ*m.

Images were again taken every 30 seconds for 15minutes to visualize immune cell activity. Cell migration in the tibia marrow cavity was distinct compared to the MT3 (Supplemental Video S3). EGFP labelled immune cell signal was very weak with no noticeable movement. However, a small number of THG-positive cells were observed migrating short distances (Figure 14).

**Fig. 14.**
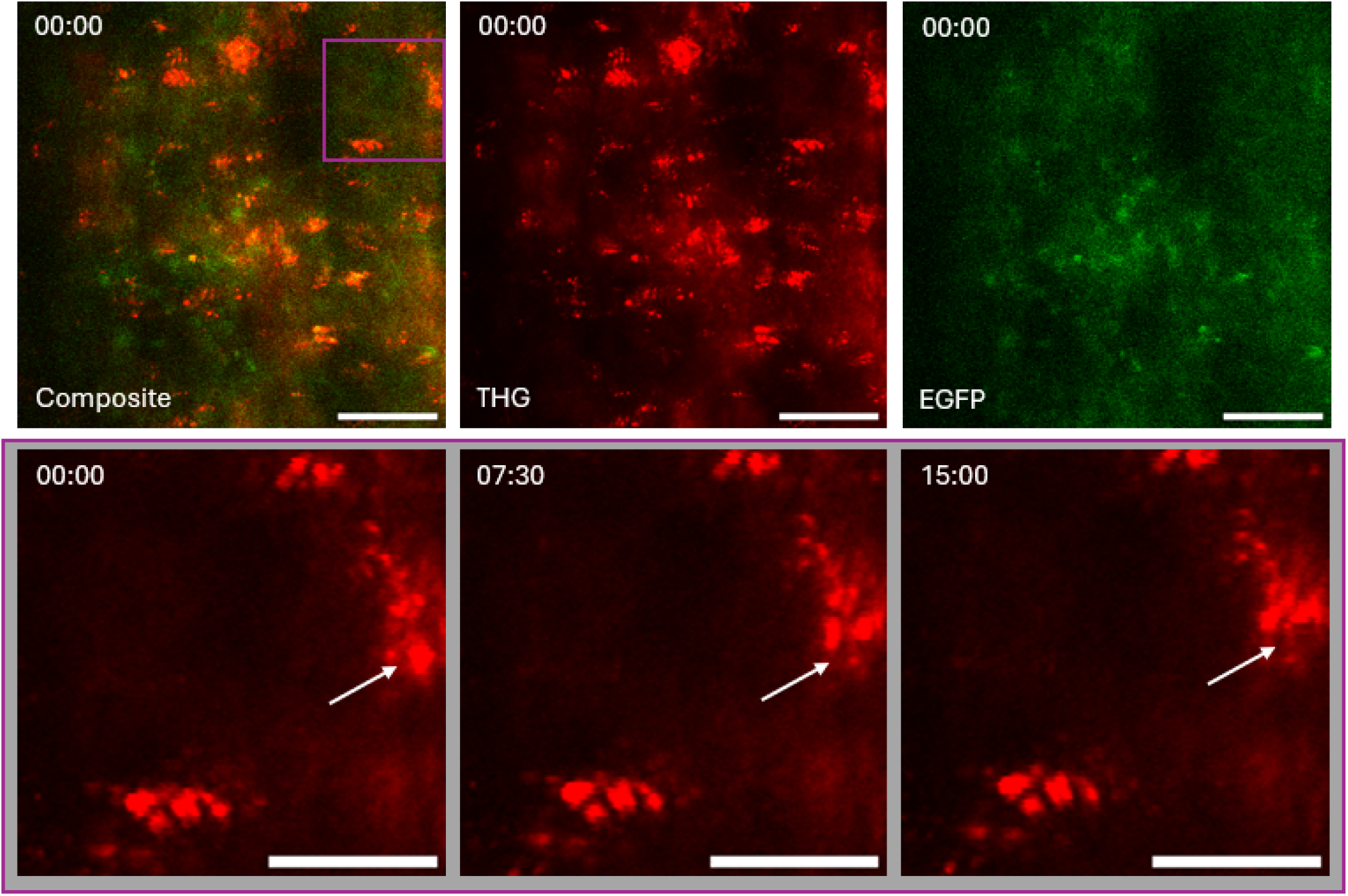
*In situ* images in the tibia marrow cavity over a 15 minute time period using 3P microscopy. (Top row) Many small, clustered cells were visible in THG (red), in contrast to few weakly labeled immune cells (green) Scale bar = 50*µ*m. (Bottom row) Over time, only a small subset of cells in THG appeared to migrate over time. Zoomed images feature a migrating cell (white arrow). Scale bar = 25*µ*m.

## Discussion

### A. Comparison of two-photon and three-photon microscopy in long bone cortices

Bone is a dynamic tissue whose functions extend far beyond mechanical support. Osteocytes embedded within cortical bone act as mechanosensors that regulate bone remodeling and adaptation while the bone marrow provides a specialized microenvironment for hematopoietic stem cells, mesenchymal stem cells, and immune cell development (1, 5, 13, 14). However, direct visualization of these cellular populations remains challenging due to the optically dense and highly scattering nature of mineralized bone. As a result, previous studies have been limited to superficial imaging depths or have relied on in vitro models, mechanically thinned bone preparations, or naturally thin bones such as the calvarium. Three-photon (3P) microscopy is a powerful nonlinear microscopy technique that has been demonstrated to successfully observe the marrow space in the mouse calvaria and the tibia through intact cortical bone. Further research from published work calls for the investigation of 3P microscopy into other long bones, the closer analysis of detection of fluorescent labeling within bone matrix, and the functionality of shorter more common excitation wavelengths. We compared the performance of 2P and 3P microscopy in collecting fluorescent signal of osteocytes over depth through intact bone. We found that 3P microscopy penetrates tissue better than 2P microscopy in all bones, as indicated by the effective attenuation length (EAL). Although 2P performed better than expected within the initial depth within the cortical bone, 3P produced a better image deep below the surface toward the endocortical surface.

Imaging osteocytes within their native environment of mineralized bone is difficult. Recent work detailing the full penetration of intact cortical bone in the tibia *in vivo* utilized middle infrared excitation (>1600nm) and high pulse rates (>2 MHz) (21). Results shown here indicate good images can be obtained using a lower pulse rate (1MHz) and shorter wavelength (1320nm) more amenable to common fluorophores (GFP).

Effective attenuation length (EAL) was greater in 3P and 3P+AO image stacks compared to 2P image stacks, indicating the excitation beam penetrates farther through the material before significant power loss and corresponding reduction in fluorescent intensity. This trend was consistent with expectations, as longer excitation wavelengths undergo less scattering as they propagate through tissue (24). Laser attenuation was not affected, however, by the addition of wavefront correction in the form of AO or RI-matching adhesive. This likely reflects the fact that EAL is primarily governed by the intrinsic scattering and absorption properties of the material (24). Because the structural and compositional properties of bone remain unchanged by these corrections, its bulk optical properties, and thus degree of attenuation, are preserved.

Slight variation in EAL was observed across the mouse long bones assessed here. These differences are likely driven by variations in cortical bone composition and microstructure, including collagen organization and mineralization. Bone exhibits hierarchical organization spanning the macro- to nanoscale, and mechanical load may further drive differences between bones by shaping tissue architecture in response to distinct strain environments and distributions, even within the same bone. Water content varies across cortical and trabecular bone in the femur and likely contributes to varying degrees of absorption (38). Additionally, the femur and tibia have been reported to have differing degrees of mineralization and water content (39).

Wavefront correction with AO did not generally improve 3P image quality in our experiments. This outcome may be partially attributed to a lower achievable laser power in the AO configuration of our microscope setup. Maximum surface laser power peaked at ~70mW at the bone surface when using AO, much lower than the maximum power achieved with 2P (~350 mW) or 3P (~265mW). The resulting reduced excitation power at greater imaging depths may have contributed to an apparent reduction in image quality. Further optimization of the optical path may enable higher delivered laser power and improved performance in future implementations. Nevertheless, these results indicate 3P with AO can achieve image quality comparable to conventional 3P imaging, even when operating at a lower excitation power. This is most likely partially aided by conservation of the beam area with the use of AO, as observed in computational simulations; thus, maximizing laser intensity in relation to 3P alone, especially in bones with high surface curvature such as the MT3 and humerus.

The application of RI-glue tended to increase the decay rate of both SNR and SBR in 3P+AO images. This effect may be due to additional absorption of an already weaker excitation laser by the RI-matched adhesive itself. Furthermore, chemical fixation may alter the RI of bone tissue relative to that of fresh bone, as fixation of tissue involves protein cross-linking and structural modification of the extracellular matrix. Indeed, fixation with formalin has been noted to alter mechanical properties of bone (40, 41).

Monitoring for *in vivo* markers of cell stress suggests the use of 3P microscopy for intravital imaging does not damage cells. A small number of osteocytes *in vivo* display spontaneous calcium signals (28). Here, osteocytes exposed to intermittent 3 minute bouts of laser exposure showed a temporal pattern in the proportion of spontaneously signaling cells that was comparable to that of the negative control group. Across all groups, the normalized magnitude of the spontaneous signals remained low. Additionally, endogenous fluorescence from metabolic proteins, including NADH, can serve as a proxy for cellular metabolic state, with increased accumulation previously associated with mitochondrial stress (29, 30). Osteocytes subjected to intermittent exposure reached peak levels of autofluorescence-positive cells and signal intensity similar to those observed in the negative control group. In contrast, continuously exposed osteocytes exhibited a markedly higher proportion of autofluorescence-positive cells and increased signal-to-background ratio, suggesting a stronger metabolic stress response under prolonged exposure conditions. Results here suggest intermittent imaging is well tolerated by osteocytes, with only a potential mild negative effect associated with extended continuous imaging. Future work would benefit from increased replicates, mechanical loading to assess impacts to osteocyte function under stress, and immunohistochemistry for the presence of heat-shock proteins in response to tissue-heating.

### B. Direct observation of immune cells in disparate marrow environments through intact cortical bone using 3P microscopy

Bone marrow is critical to hematopoiesis, immune cell formation, and continued support of bone cells. However, direct *in vivo* observation of marrow cell dynamics has been limited due to its location within the interior cavity of optically dense mineralized bone. The advantage observed here of increased image quality deeper within bone can be extended further toward visualization of the marrow cavity. Using 3P microscopy, we successfully observed the migration of immune cells through intact cortical bone.

The long bones examined here were noted to have a differing appearance of the marrow cavity, with the MT3 diaphysis containing numerous large adipocytes not seen in the tibia metaphysis. This distribution reflects the natural transition observed with age from red to yellow marrow in the diaphysis of the extremities and long bones, with the metaphyses affected last while the epiphyses retain red marrow (42, 43). Thus, while only one region of the MT3 and tibia was examined here, a single bone may still potentially serve as a strong model for the study of both red and yellow marrow by capitalizing on the marrow distribution in different regions of long bones. This would enable direct comparison of hematopoietic and mesenchymal marrow niches under shared physiological and structural conditions.

The MT3 and tibia also maintained a stark contrast in EGFP labelled immune cells. Only a few weakly labeled immune cells were observed in the marrow cavity of the tibia in contrast to the robust labeling observed in the MT3. One possible explanation for this low signal is heterogeneity in CX3CR1 expression, the promoter gene used for fluorescent labeling in our experiment. The CX3CR1 gene is broadly expressed in many immune cells from the hematopoietic lineage, but expression level varies with maturation state (44). A potential alternative and complimentary factor may be limited excitation through the thicker cortical bone of the tibia paired with the restriction of substantially increasing laser power.

A cell population largely lacking GFP labeling was noted in the tibia marrow via THG, forming small clusters with subsets migrating over time. Rakhymzhan et al. observed fluorescently labeled B lineage cells in the marrow, which appeared similar in size, shape, quantity, and placement to unknown cells observed here (21). Additionally, they found an inverse link between THG intensity and migratory behavior, aligning with our observation of few conspicuous cells moving over time. Their findings point to the cells observed here as being plasma cells as well. Although B cells largely do not express CX3CR1, a small subset of CX3CR1+ B cells has been reported in human blood and may account for slight overlap with EGFP signal (45). Regardless, THG may also be used to effectively observe cells in the marrow cavity without fluorescent labeling.

Imaging of the tibia was not performed *in vivo* in this study. Abrupt loss of blood supply and pressure may negatively impact cells within the marrow space, skewing the brightness of fluorescent markers and migration behaviors observed here. *In vivo* imaging is an important follow-up and may build on established methodologies. Prior studies have facilitated tibial imaging through removal of overlying soft tissue and stabilization of the lower hindlimb by securely clamping the ankle and knee (21, 46). Thus, *in vivo* imaging of the tibia could be achieved for future studies.

Altogether, our results demonstrate that 3P microscopy consistently outperforms 2P microscopy for deep bone imaging as well as supporting the continued utility of 2P microscopy for superficial bone imaging, particularly when enhanced with an RI-matched adhesive. We further demonstrated the capability of 3P microscopy to image through intact cortical bone and into the marrow cavity, enabling successful observation of immune cell migration *in vivo*. Pairing 3P microscopy with intravital imaging of the MT3 provides a promising new model for studying marrow cell populations, while imaging in the tibia offers complementary opportunities to investigate red marrow environments. Collectively, this work expands the potential for studies of osteocytes, marrow cell populations, and their interactions within the native bone microenvironment.

## ACKNOWLEDGEMENTS

We thank the Schaffer-Nishimura group at Cornell for providing access to their three-photon microscope. We are especially grateful to Nicole Chernavsky for training in the use of the microscope and ongoing technical support and troubleshooting. We also thank Julia Horn and Anthea Spirko for their assistance in developing a data analysis workflow for evaluating image quality from image stacks.

## Supplemental

### Normal intensity and SNR of fluorescent signal through cortical bone

The normal intensity of fluorescent GCaMP8m signal was comparable to or lower with 2P than with 3P in all bones (Figure S1). In the MT3, 2P produced the lowest signal, although it decayed at the same rate as 3P, while 3P+AO did not perform better than 3P alone. Adding RI-matched glue improved the 2P decay rate but lowered the peak normal intensity. In the tibia, RI-matched glue improved 2P image intensity to 3P levels and improved the 3P+AO decay rate. In the femur, 3P showed the slowest loss of normal intensity with depth. The most dramatic effect of adding RI-matched glue was observed in the femur, where decay rates improved for 2P, 3P, and 3P+AO. Image stacks collected using 3P showed relatively consistent normal intensity, with a slight increase in some cases with depth.

Trends in normal intensity were not reproduced when evaluating the signal-to-noise ratio (SNR) (Figure S2). Surprisingly, 2P generated higher SNR within approximately 50–80 *µ*m of the surface. However, its SNR decreased rapidly with depth and eventually declined below that of 3P and 3P+AO. Consistent with the normalized intensity results, 3P+AO did not perform better than 3P alone. Adding RI-matched glue generally increased peak SNR values, but potential improvements with depth were offset by faster SNR decay in 3P and 3P+AO images. Interestingly, RI-matched glue extended the depth range over which 2P SNR remained higher than 3P and 3P+AO by approximately 10–20 *µ*m. Despite the pronounced effect of RI-matched glue on normalized intensity in the femur, it did not produce a measurable change in SNR.

**Fig. S1.**
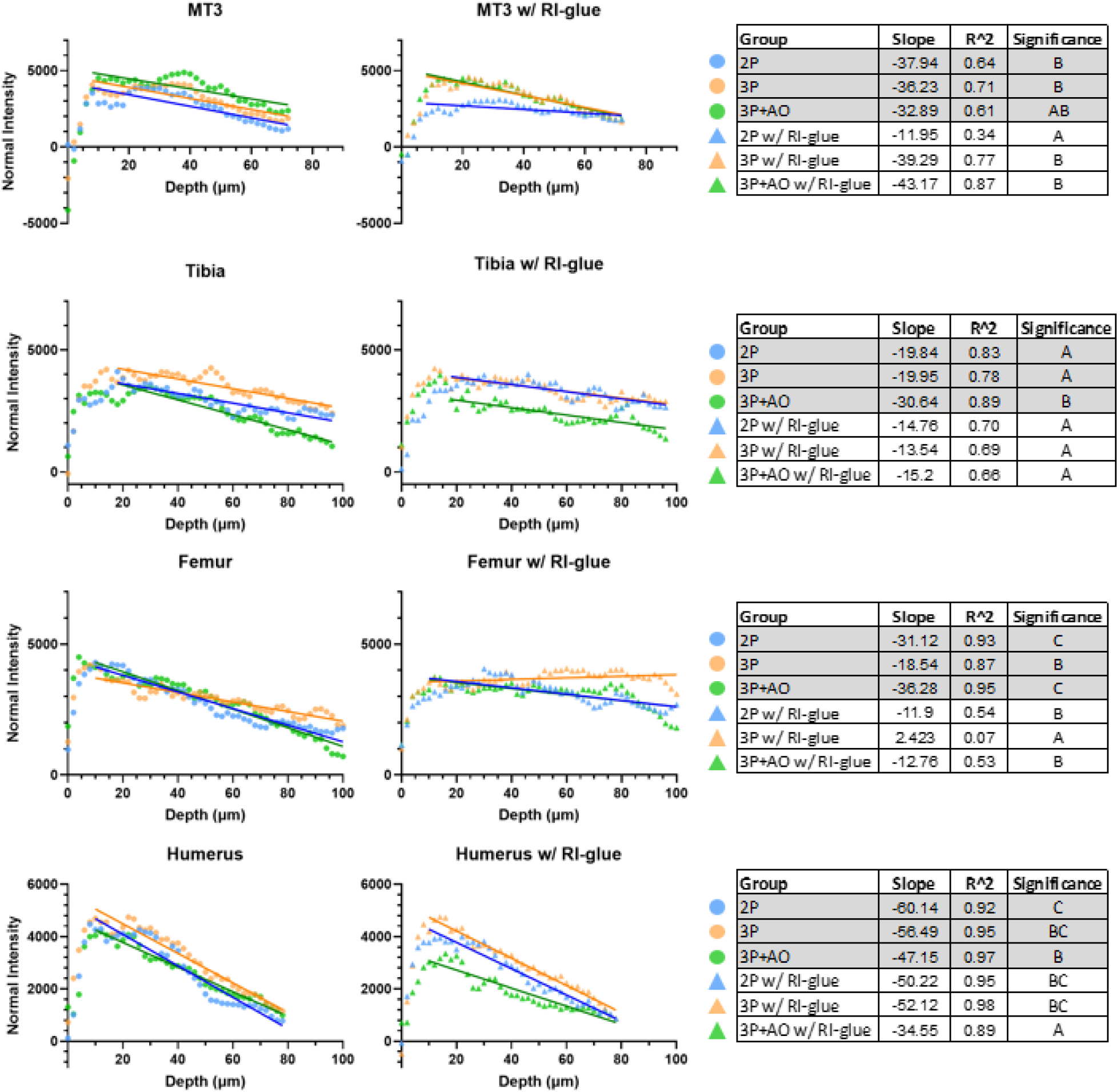
Normalized intensity of osteocyte GCaMP8m signal over depth in 2P and 3P microscopy. Images collected using 2P were generally comparable to those obtained with 3P microscopy. The addition of refractive index (RI)-matched glue was most effective in the femur, substantially reducing the decay in normalized intensity with imaging depth across 2P, 3P, and 3P+AO image stacks.

**Fig. S2.**
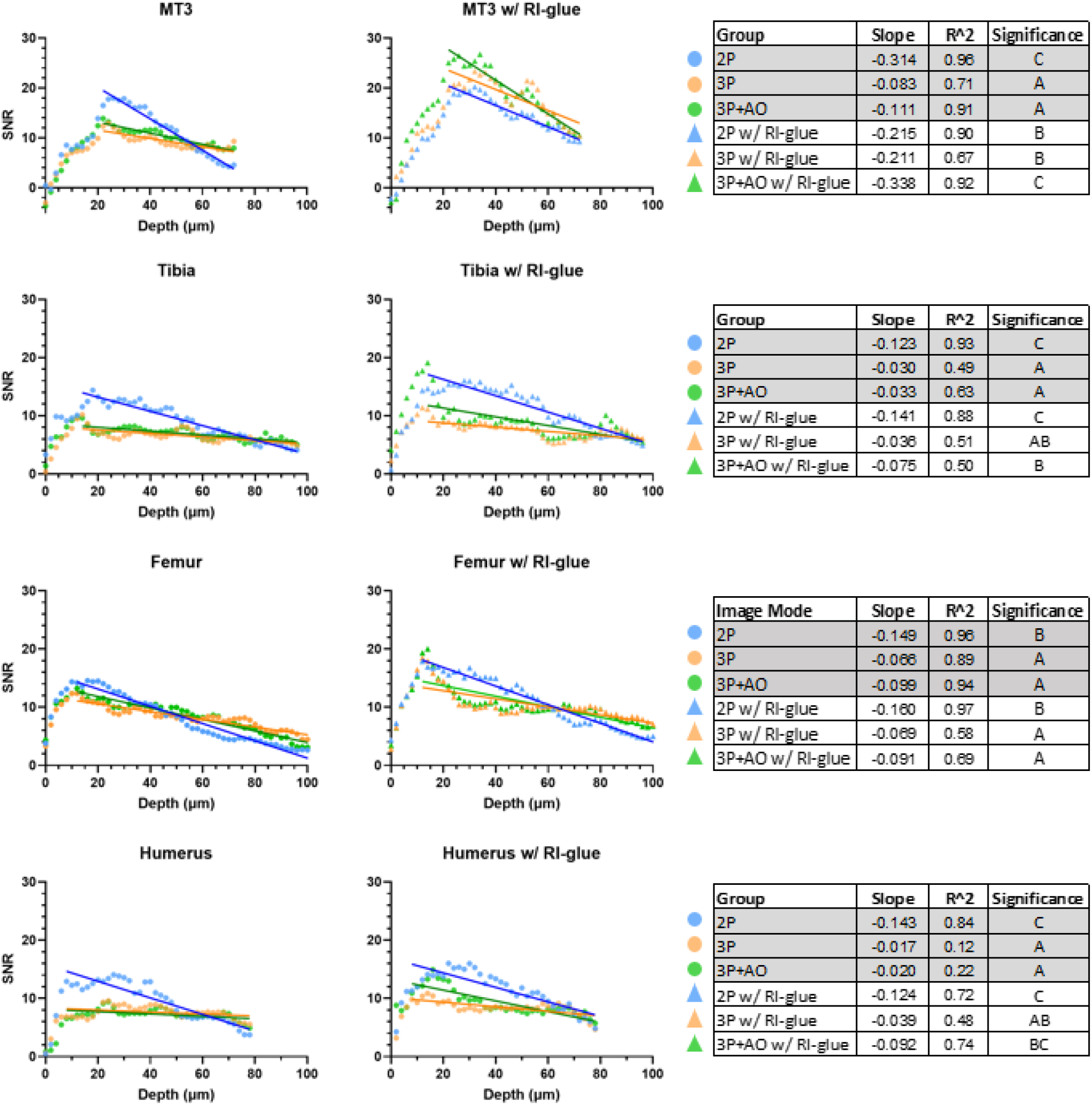
Signal-to-noise ratio (SNR) of osteocyte GCaMP8m signal over depth in 2P and 3P microscopy. Two-photon microscopy generated higher SNR to approximately 50–80 *µ*m below the surface, and RI-matched glue extended the depth over which 2P SNR remained higher. RI-matched glue often increased SNR decay in 3P and 3P+AO image stacks.

**Fig. S3.**
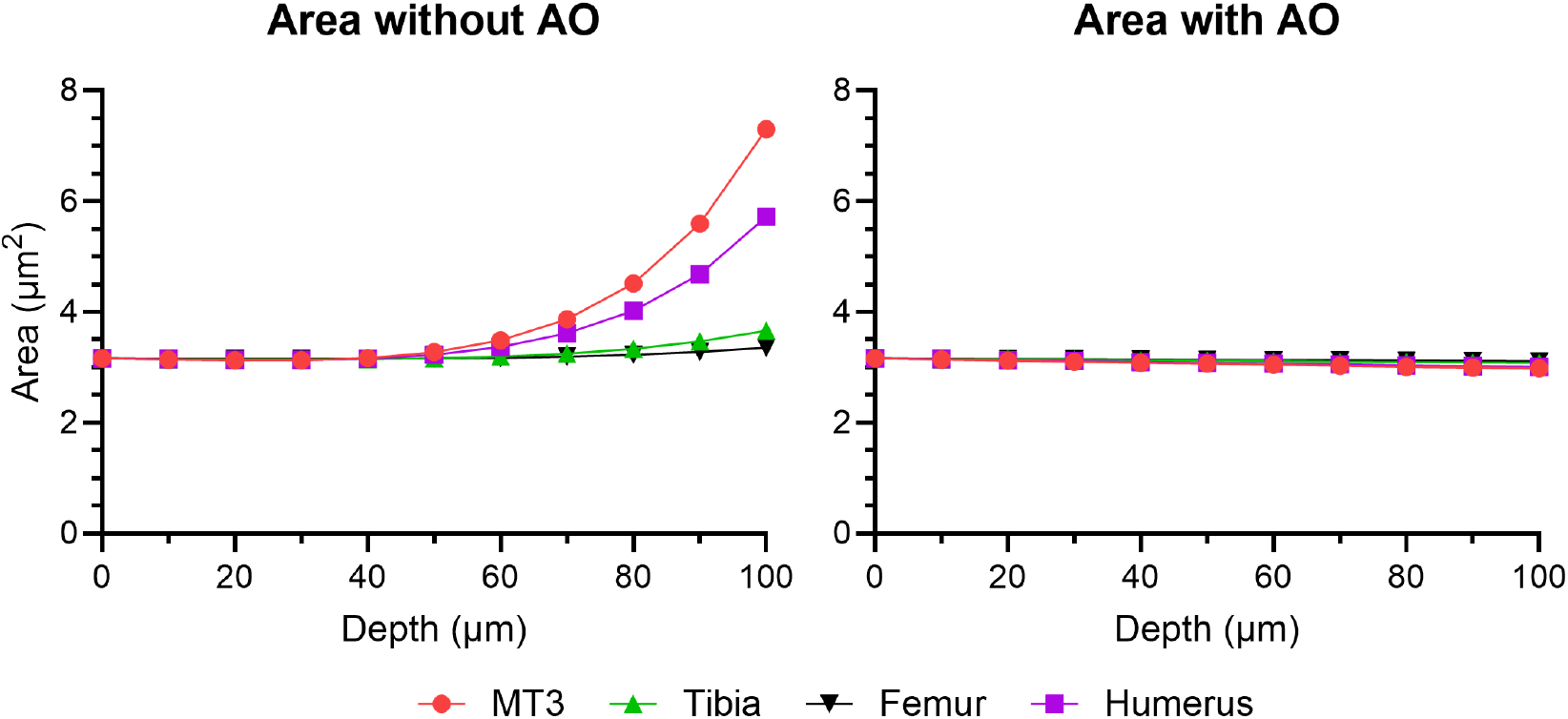
Simulation of beam area with imaging depth beneath a curved surface. Without adaptive optics, beam area increased exponentially with depth, particularly in bones with high surface curvature such as the MT3. Adaptive optics effectively maintained beam area with depth.

**Fig. S4.**
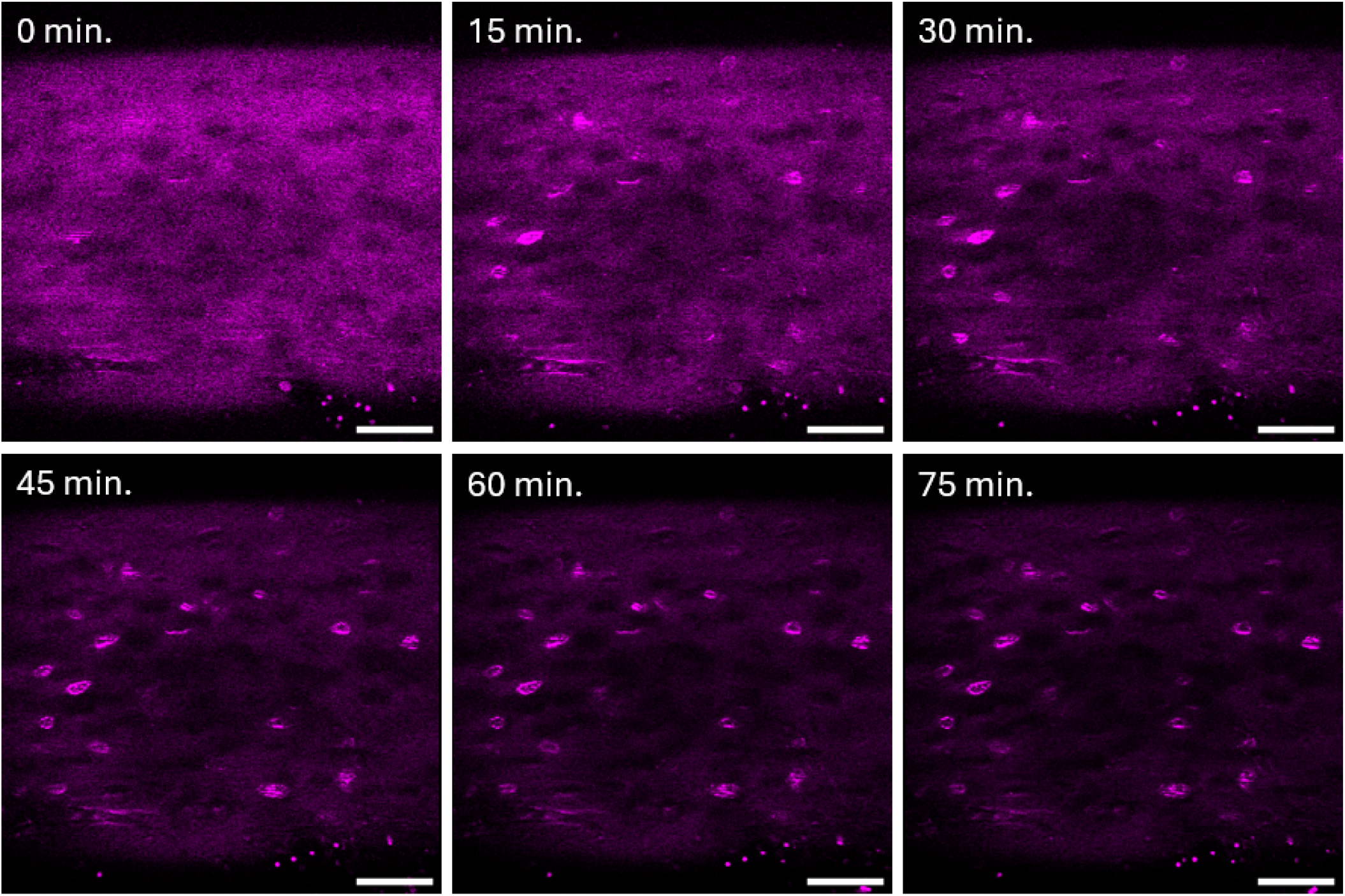
Representative images depicting the increase in autofluorescence over time following intermittent laser exposure in a mouse MT3. Scale bar=50*µ*m.

**Table S1.** Amplitude of simulated contributing Zernike polynomials as imaging depth increases at the apex of the surface curvature.

| Depth ($\mu\text{m}$) | Primary Vertical Astigmatism (Z6) | | | | Primary Spherical (Z11) | | | |
| --- | --- | --- | --- | --- | --- | --- | --- | --- |
|  | MT3 | Tibia | Femur | Humerus | MT3 | Tibia | Femur | Humerus |
| 0 | 0.00 | 0.00 | 0.00 | 0.00 | 0.00 | 0.00 | 0.00 | 0.00 |
| 10 | 0.00 | 0.00 | 0.00 | 0.00 | 0.00 | 0.00 | 0.00 | 0.00 |
| 20 | 0.01 | 0.00 | 0.00 | 0.00 | 0.00 | 0.00 | 0.00 | 0.00 |
| 30 | 0.01 | 0.00 | 0.00 | 0.01 | 0.00 | 0.00 | 0.00 | 0.00 |
| 40 | 0.02 | 0.01 | 0.01 | 0.02 | 0.00 | -0.01 | 0.00 | 0.00 |
| 50 | 0.03 | 0.01 | 0.01 | 0.03 | -0.01 | -0.01 | -0.01 | -0.01 |
| 60 | 0.05 | 0.02 | 0.01 | 0.04 | -0.01 | -0.01 | -0.01 | -0.01 |
| 70 | 0.06 | 0.02 | 0.02 | 0.05 | -0.01 | -0.01 | -0.01 | -0.01 |
| 80 | 0.08 | 0.03 | 0.02 | 0.07 | -0.01 | -0.01 | -0.01 | -0.01 |
| 90 | 0.10 | 0.04 | 0.03 | 0.09 | -0.01 | -0.01 | -0.01 | -0.01 |
| 100 | 0.13 | 0.05 | 0.03 | 0.11 | -0.01 | -0.01 | -0.01 | -0.01 |

**Video S1**. Representative video of spontaneous calcium signaling in the mouse MT3. One cell displaying a calcium transient is highlighted by a white arrow.

**Video S2**. Time-lapse imaging over 15 min showing cell migration in the marrow of a mouse MT3 diaphysis. Scale bar = 20 *µ*m.

**Video S3**. Time-lapse imaging over 15 min showing cell migration in the marrow of a mouse tibial metaphysis. Scale bar = 20 *µ*m.

