## Supplementary figures and images for "Depth-Dependent Advantages of Three-Photon Microscopy for Imaging Through Intact Murine Cortical Bone and Into the Marrow"

### Supplemental Video 1

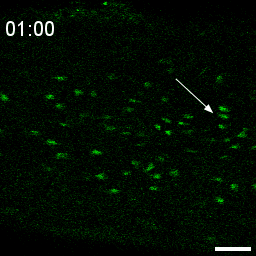

### Supplemental Video 2

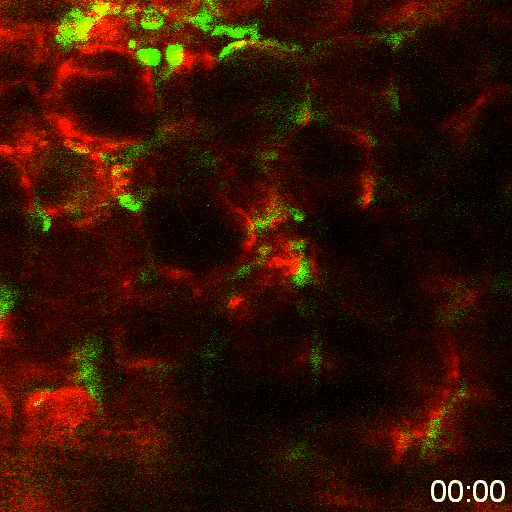

### Supplemental Video 3

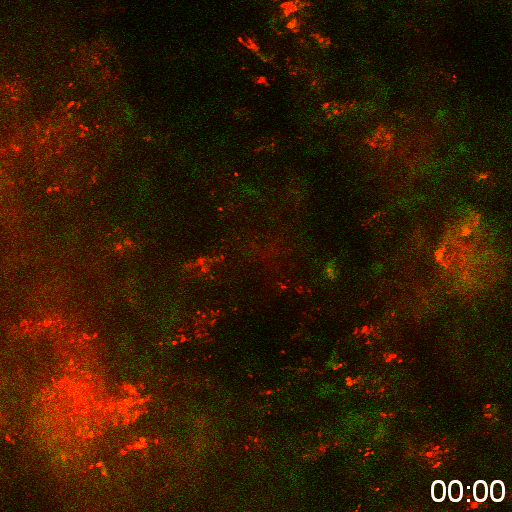
